# Spatiotemporal regulation of arbuscular mycorrhizal symbiosis at cellular resolution

**DOI:** 10.1101/2025.10.31.685811

**Authors:** Tania Chancellor, Gabriel Ferreras Garrucho, Garo Z. Akmakjian, Héctor Montero, Sarah Bowden, Matthew S. Hope, Emma Wallington, Samik Bhattacharya, Christian Korfhage, Julia Bailey-Serres, Uta Paszkowski

## Abstract

Arbuscular mycorrhizal (AM) symbiosis develops through successive colonization of root epidermal and cortical cells, culminating in the formation of arbuscules, tree-like intracellular structures that are transient yet essential sites of nutrient exchange. To dissect the cellular and structural complexity of AM establishment in rice roots colonized by *Rhizophagus irregularis*, we applied dual-species spatial transcriptomics to simultaneously monitor plant and fungal gene transcripts at single-cell resolution. This approach revealed surprising differences in transcriptional activity between fungal structures and showed that morphologically similar arbuscules can be transcriptionally distinct. These findings suggest hidden functional diversity among arbuscules at single-cell resolution. Because arbuscules form and degenerate within only a few days, we further sought to capture translational activities across their life span. We pioneered AM-inducible TRAP-seq (Translating Ribosome Affinity Purification followed by RNA-seq) using stage-specific promoters, enabling cell-type- and stage-resolved profiling in AM symbiosis. This revealed extensive spatiotemporal reprogramming of nutrient transport and signalling, with distinct sets of phosphate, nitrogen, and carbon transporters and regulators induced or repressed at different stages of arbuscule development, suggesting that nutrient exchange is dynamically regulated across the arbuscule life cycle. More broadly, cell wall biosynthesis genes and key defence markers were suppressed during arbuscule formation, whereas at a later stage, defence markers were strongly upregulated, suggesting a host-driven shift towards arbuscule termination. Together, these findings highlight the nuanced and dynamic regulation of AM symbiosis at the cellular level, refining our understanding of how nutrient exchange and fungal development are coordinated in space and time.

## 2. Introduction

The symbiotic relationship between plants and arbuscular mycorrhizal (AM) fungi is both ancient and widespread, with over 80% of extant plant species engaging with AM fungi across diverse ecosystems, including domesticated crop species in agricultural settings. The relationship is based on a finely balanced nutritional mutualism. AM fungi greatly increase the uptake of mineral nutrients such as phosphate and nitrogen, whereas the plant provides the fungus with essential carbon in the form of fatty acids, necessary for the completion of the AM fungal life-cycle (Bravo et al., 2017; Jiang et al., 2017; Keymer et al., 2017; Luginbuehl et al., 2017).

The establishment of AM symbiosis is tightly coordinated and begins with mutual recognition, followed by the fungus contacting the host root, and initiating a complex reprogramming of plant cellular architecture that facilitates fungal entry and colonization of the root epidermal and cortical cell layers. Central to the symbiosis is the formation of arbuscules, finely branched fungal structures within root cortical cells. Arbuscules have a short lifespan, forming and collapsing within a few days. They are enveloped by the plant-derived peri-arbuscular membrane (PAM), which is enriched in specialized transporters mediating the bidirectional flow of minerals and fatty acids that underpin the mutualistic interaction. Following arbuscule development, the fungus accumulates host-derived lipids and, in some AM clades, forms storage vesicles inside the root cortex, while reproductive spores develop both inside and outside the root (figure 1A) (Smith & Read, 2010).

**Figure 1.**
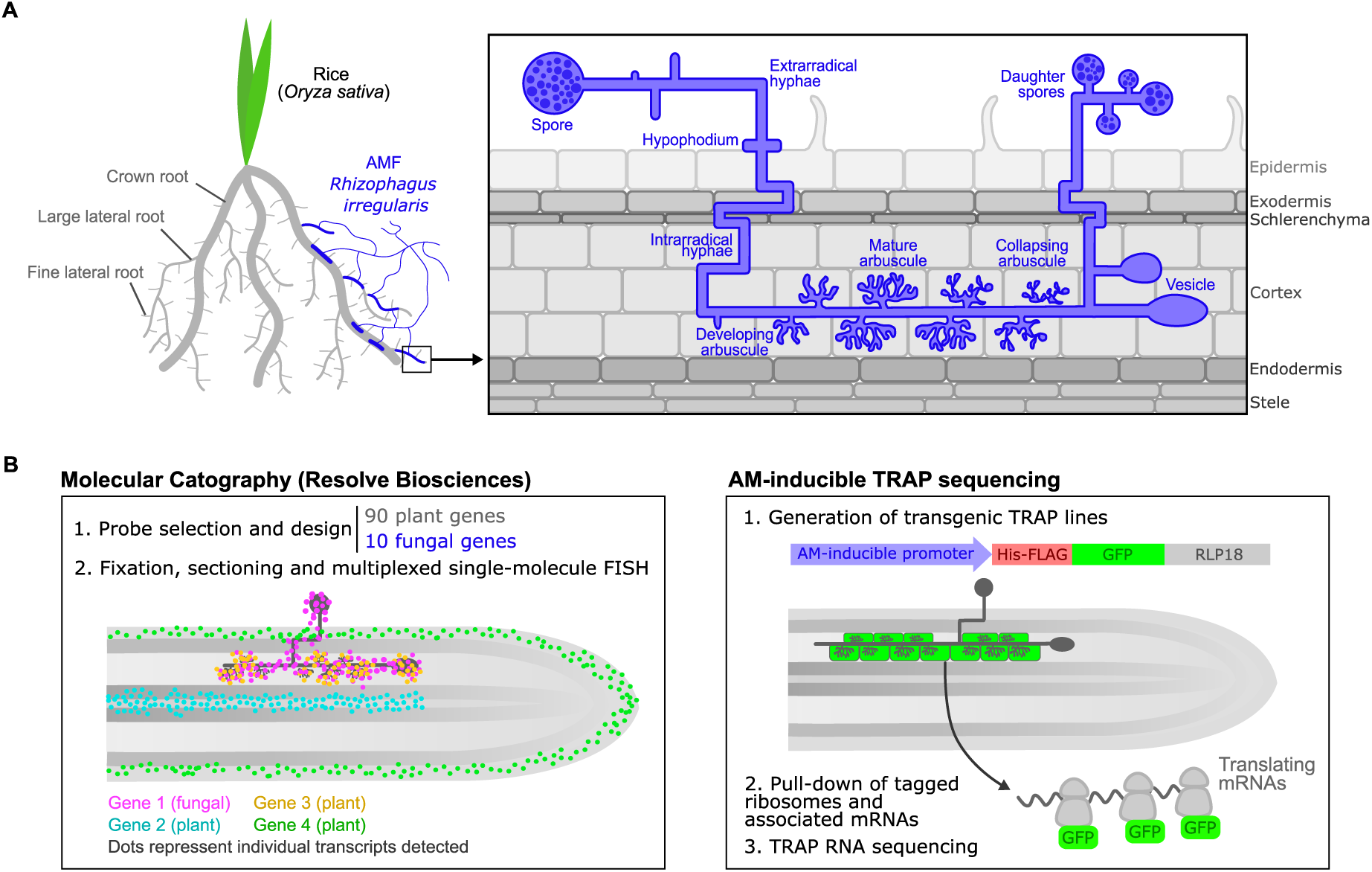
Graphical abstract of the spatiotemporal study of arbuscular mycorrhizal symbiosis between rice and *Rhizophagus irregularis*. (A) Simplified representation of the biological system of study, the mutualism between roots of rice (*Oryza sativa*) and the arbuscular mycorrhizal fungus (AMF) *R. irregularis*, with the macroscopic view of the interaction on the left, and the microscopic view of symbiotic fungal structures outside and within the plant root on the right. (B) Schematic representation of the high-resolution experimental techniques employed in this study to resolve the spatiotemporal dynamics of AM symbiosis. On the left, Molecular Cartography by Resolve Biosciences, including a simplified mycorrhizal root section with 4 genes detected. FISH, Fluorescence in-situ Hybridization. On the right, Translating-Ribosome Affinity Purification (TRAP) RNA sequencing using AM-inducible promoters, indicating the structure of the TRAP construct (with a 6x-His-FLAG as well as GFP tags fused to the ribosomal protein RLP18) and hypothetical expression pattern of the construct based on AM-inducible promoter used.

The dynamic and asynchronous nature of fungal root colonization, with co-occurring distinct fungal structures in adjacent plant cells, poses challenges to the study of discrete stages of the association. Similarly, molecular mechanisms that act cell-autonomously to coordinate the rapid formation and turnover of arbuscules remain difficult to identify. As a result, little is known regarding the fine-scale spatiotemporal regulation of AM development. Though vital spatial information has been gained from the use of transcriptional and translational reporters, RNA in-situ hybridization, and laser microdissection coupled with RNA-seq, such methods fail to capture the overall complexity of the symbiosis. Recent advances such as spatial transcriptomics have expanded our ability to interrogate spatial gene expression in AM symbiosis (Ferreras-Garrucho et al., 2025). Using single-nucleus sequencing and spatial transcriptomics (Visium, 10X Genomics), Serrano et al. (2024) identified coordinated, stage-specific transcript abundance programs during the symbiosis between *Medicago truncatula* and *Rhizophagus irregularis*. However, insights from Visium spatial transcriptomics were constrained by limited resolution, with each capture spot encompassing transcripts from approximately 1-5 plant cells, thus limiting the ability to resolve cell-type-specific or structure-specific transcriptional programs.

In this study, we combined high-resolution spatial transcriptomics with Translating Ribosome Affinity Purification and RNA sequencing (TRAP-seq) to conduct a detailed dissection of the spatial and temporal regulation of AM symbiosis between rice (*Oryza sativa*) and *R. irregularis* (figure 1B). Using the Molecular Cartography™ spatial transcriptomics platform by Resolve Biosciences (a combinatorial smFISH based approach), we targeted a panel of 100 key AM symbiosis genes (90 plant and 10 fungal targets), providing the first spatially defined, dual-species transcript pattern resource at single-cell resolution. We then leveraged TRAP-seq to provide a complementary insight into translational reprogramming across AM development (Reynoso et al., 2015, 2022; Zanetti et al., 2005; Zhao et al., 2017). Here, we selected three rice promoters known to be activated at different stages of AM colonization. Our AM-inducible TRAP-seq approach enabled targeted investigations of gene activity at distinct developmental time frames. The high-resolution data presented here provides new insights into the spatial expression of key plant and fungal AM genes, uncovers significant spatiotemporal repatterning of key nutrient transport and signalling genes, and sheds new light on translational reprogramming from arbuscule development through to arbuscule degeneration. Together, our data contribute to an improved understanding of the fine-scale regulation of AM symbiosis in rice.

## 3. Results

### 3.1 Dual-species spatial transcriptomics at single-cell resolution

To understand how gene regulation supports the dynamic progression of arbuscular mycorrhizal (AM) symbiosis, we investigated the spatial expression of key symbiotic genes across different stages of root colonization. Here, we curated a panel of 100 targets from rice and the model arbuscular mycorrhizal fungus *Rhizophagus irregularis*, comprising 10 fungal genes associated with AM symbiosis and 90 plant genes. The plant set included 25 genes involved in AM signalling, 22 established cell type–specific markers, and genes implicated in the perception, transport, and signalling of nitrogen (11 genes), phosphate (20 genes), and carbon (12 genes). To capture fungal structures across AM development, we compared noncolonized roots (−Ri) with *R. irregularis* colonized roots (+Ri) harvested at 6 weeks post inoculation (wpi). Two experimental replicates were performed to strengthen detection of transcript patterns across colonization stages. To capture spatial gene expression across the complex rice root system, our spatial transcriptomics experiments included both crown roots (CRs) and large lateral roots (LLRs), which are transcriptionally distinct in the noncolonized state but based on bulk mRNA-seq data, exhibit convergent transcript profiles upon arbuscular mycorrhizal colonization (Gutjahr, Sawers, et al., 2015).

Following 100-plex RNA fluorescent *in situ* hybridization (smFISH, Molecular Cartography, Resolve Biosciences), probe performance and section quality were assessed, and samples/probes were filtered before proceeding with data analyses. Probes with less than an average of 20 transcripts per section in both noncolonized and mycorrhizal conditions were removed from further analysis, leaving a total of 74 probes (70 plant, 4 fungal). Of 30 sections, 20 were of sufficiently high-quality, leaving a total of 6 sections from experiment 1 (2 noncolonized, 4 mycorrhizal), and 14 sections from experiment 2 (6 noncolonized, 8 mycorrhizal). To scrutinize transcript abundances across all samples, we generated a gene count matrix from the transcript spot data. Principal Component Analysis (PCA) revealed a distinct separation of samples based on mycorrhizal status along PC1, which accounted for 68% of the variance. Notably, PC2 captured a modest variation (9%) within mycorrhizal samples across the two experiments (figure S1A), likely reflecting differences in capture rates of 24 low-abundance transcripts (figure S1B). Transcript abundance did not differ significantly between CR and LLR root-types except for two genes (the vascular markers *Os01g0803300* and *OsUMAMIT9*), whereas more than half of the set, 48 out of 74 genes, were differentially abundant between *R. irregularis*-colonised and mock samples (figure S2). To focus on +Ri vs −Ri differences and to maximise the range of cell-types captured, CR and LLR samples have been considered together for the remainder of the study, unless otherwise mentioned.

Composite imaging with DAPI and WGA-AF488 revealed a clear blueprint of plant and fungal organization, highlighting nuclei and AM fungal structures, respectively. DAPI staining confirmed earlier observations (Bianciotto et al., 1995) that fungal nuclei are absent from fine arbuscule branches, but present in the arbuscule trunk and intraradical hyphae. Fungal nuclei were also abundant in vesicles (figure 2A). To validate the dual-species Molecular Cartography approach, we assessed the spatial expression of cell-type marker genes across all sections. A total of 22 cell-type marker probes were selected from a list validated by (Zhu et al., 2025) based on published single-cell datasets in rice (Liu et al., 2021; Wang et al., 2021; Zhang et al., 2021). Of these, 15 markers yielded sufficient transcript counts (>20 transcripts per section) and were carried forward for detailed analysis. In noncolonized samples, 10 of the 15 markers exhibited spatial expression patterns consistent with their expected tissue localization. For example, the exodermal marker *GDSL esterase/lipase protein 7* (*OsGELP7*), the cortical marker *Redox Associated Intermediate 1* (*OsRAI1*), and two uncharacterized phloem-enriched translated mRNAs (*Os07g0634400* and *Os01g0803300*) (Reynoso et al., 2022) aligned closely with their predicted distributions (figure 2B). In contrast, the phloem-expressed *Amino Acid Permease 11G* (*OsAAP11G*), the atrichoblast-expressed nitrate transporter *OsNRT2.3* and the endodermal marker *Os10g0459300* displayed nonspecific spatial expression patterns (figure S3). Their diverging expression patterns from previous spatial and single-cell transcriptomics datasets (Liu et al., 2021; Wang et al., 2021; Zhang et al., 2021; Zhu et al., 2025) might be due to the distinct sample type. Compared to embryonic root tips grown in hydroponics or plates, we sampled the mature zone of non-embryonic roots grown in sand. This highlights the importance of identifying robust cell-type markers across developmental ages, root zones, root types and substrates.

**Figure 2.**
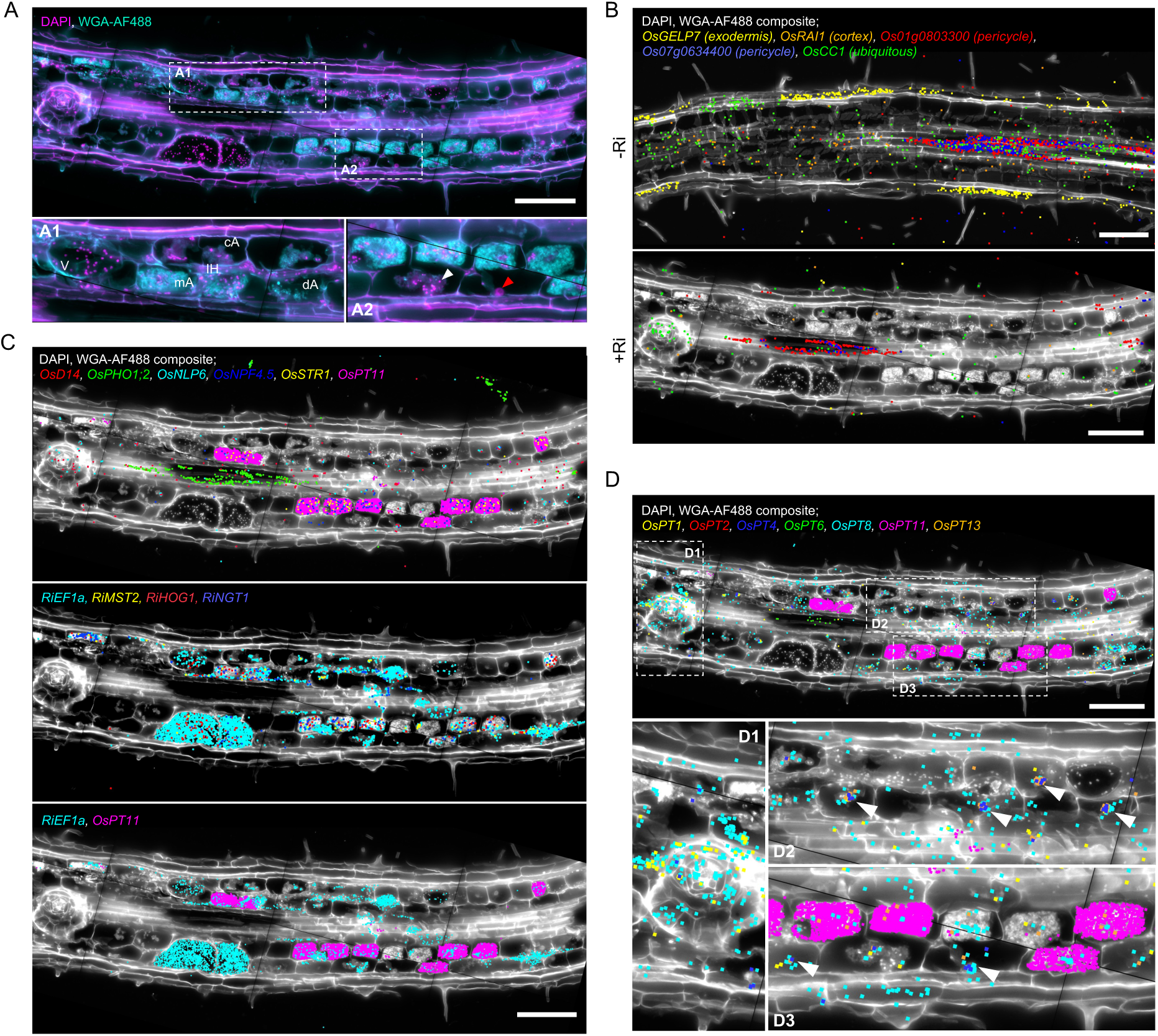
Molecular Cartography for dual species spatial transcriptomics of arbuscular mycorrhizal symbiosis between rice and *R. irregularis*. (A) Composite image of DAPI (magenta) and WGA-AF488 (cyan) of an *R. irregularis*-inoculated (+Ri) section for visualization of nuclei, cell boundaries and fungal structures. Close-up A1 showcases all fungal structures visualized: intraradical hyphae (IH), arbuscules at distinct development stages (developing, dA; mature, mA; and collapsing, cA), and vesicles (V). Close-up A2 showcases the difference between plant (red), and fungal (white arrow) nuclei. (B) Image-transcript overlays showing spatial expression of cell-type specific markers in a −Ri and a +Ri section. (C) Image-transcript overlays comparing spatial expression of plant and fungal transcripts in a +Ri section, including key AM markers. (D) Image-transcript overlays showing spatial expression of the seven most abundant plant phosphate transporters. Close-ups D1 to D3 highlight differing spatial detection patterns in diverse cell-types: D1 in emerging lateral root, D2 and D3 for arbuscules at distinct developmental stages and non-colonised cortical cells. Arrows highlight clusters of *OsPT4*, *OsPT8* and *OsPT13* transcripts around plant nuclei of non-colonised and collapsing arbuscule-containing cells. DAPI/WGA-AF488 composite in white, different colours correspond to independent transcripts, each spot corresponds to one detected transcript. Scale bar, 100 μm.

Interestingly, the transcript abundance of most cell-type markers (11/15) was reduced in mycorrhizal samples (figure 2B, S2). When sufficiently captured, such as highly expressed *Os07g0634400* and *Os01g0803300* they retained their expected spatial patterning in colonized roots (figure 2B). Importantly, the ubiquitously expressed *Cytochrome C1* (*OsCc1*) was not downregulated, suggesting that the reduction of cell-type marker transcripts is unlikely to be a technical artefact and instead may reflect a biological consequence of AM symbiosis (figure S2). It seems therefore plausible that intraradical fungal colonisation markedly affects cell-type identity, or at least expression of cell identity markers.

Next, we compared detection of plant and fungal transcripts, particularly known markers induced during AM symbiosis, to validate the capacity of this technology for dual-species spatial transcriptomics. Differential gene expression analysis revealed significant up-regulation of all fungal genes and many known plant AM markers between mycorrhizal and non-mycorrhizal roots (figure S2). These included the nutrient transporters Phosphate Transporter 11 (*OsPT11*), Nitrate/Peptide Family Transporter 4.5 (*OsNPF4.5*), and Stunted Arbuscule 1 (*OsSTR1*), which were specifically localized to arbuscule-containing cells in mycorrhizal roots. Fungal transcripts closely tracked the distribution of fungal structures in colonized roots, and we were able to successfully capture transcripts of the two species within the bounds of a single plant cell (figure 2C, S4, S5). Notably, the fungal marker gene *RiEF1ɑ*, encoding an essential translation elongation factor, had striking transcript abundance within vesicles, where transcripts co-localized with fungal nuclei. In contrast, significantly fewer *RiEF1ɑ* transcripts were detected in arbuscules (figure 2C, S4, S6, S7). Additional fungal genes *RiHOG1, RiMST2*, and *RiNGT1*, also closely tracked vesicles, hyphae and arbuscules in similar distribution to *RiEFa*, yet with lower abundance, with *RiHOG1* likewise showing evidence of nuclear colocalization in vesicles (figure 2C, S4, S7). This nuclear association suggests that vesicles are not simply passive storage bodies but transcriptionally dynamic compartments, with active expression of certain genes concentrated around nuclear domains.

As phosphate is essential for AM symbiosis in signalling, regulation and nutrient exchange, and our panel included most high-affinity phosphate transporters in rice, a targeted investigation of their expression patterns seemed valuable. The symbiotic phosphate transporters *OsPT11* and *OsPT13* were significantly induced in mycorrhizal roots (figure S2). *OsPT11* expression was restricted to arbuscule-containing cells, whereas *OsPT13* was also detected in non-colonised cortical cells (figure 2D, S4). Interestingly, non-symbiotic PTs had distinct expression patterns between +Ri and −Ri roots. *OsPT6* was largely confined to the stele regardless of colonisation, but was significantly down-regulated in mycorrhizal roots. Conversely, *OsPT1* was significantly induced in +Ri roots, highly abundant around emerging laterals and stele (figure 2D, S2, S4, S5). *OsPT8* was the most abundant and expressed widely across most cell types in both −Ri and −Ri roots, including in arbuscule-containing cells (figure 2D, S4, S5). Furthermore, in mycorrhizal roots, *OsPT8,* as well as the less abundant *OsPT4* and *OsPT13*, showed a striking sub-cellular spatial association with plant nuclei in uncolonized cells and in cells containing collapsing arbuscules (figure 2D, S4). Together, these patterns highlight diverse spatial domains and colonisation-dependent regulation of phosphate transporters, extending beyond a simple symbiotic versus non-symbiotic division.

### 3.2 Single-cell clustering and colocalization analyses

To investigate the spatial integration of AM symbiosis genes, we calculated transcript colocalization scores for all 74 targets, excluding low-abundance genes that showed poor self-colocalization (score < 0.2). Despite transcript counts being generally low in noncolonized samples, four distinct clusters emerged, largely reflecting rice cell types (exodermis, vascular tissue, xylem/phloem, and cortex; figure S8). By contrast, colonized samples displayed a markedly altered transcriptional landscape, with five distinct clusters that reflected a profound reorganization of spatial expression patterns across root tissues (figure 3A, B). Spatial colocalization patterns were broadly consistent between CR and LLR root types, indicating no clear differences in the spatial organization of AM symbiosis genes between these root types (figure S8).

**Figure 3.**
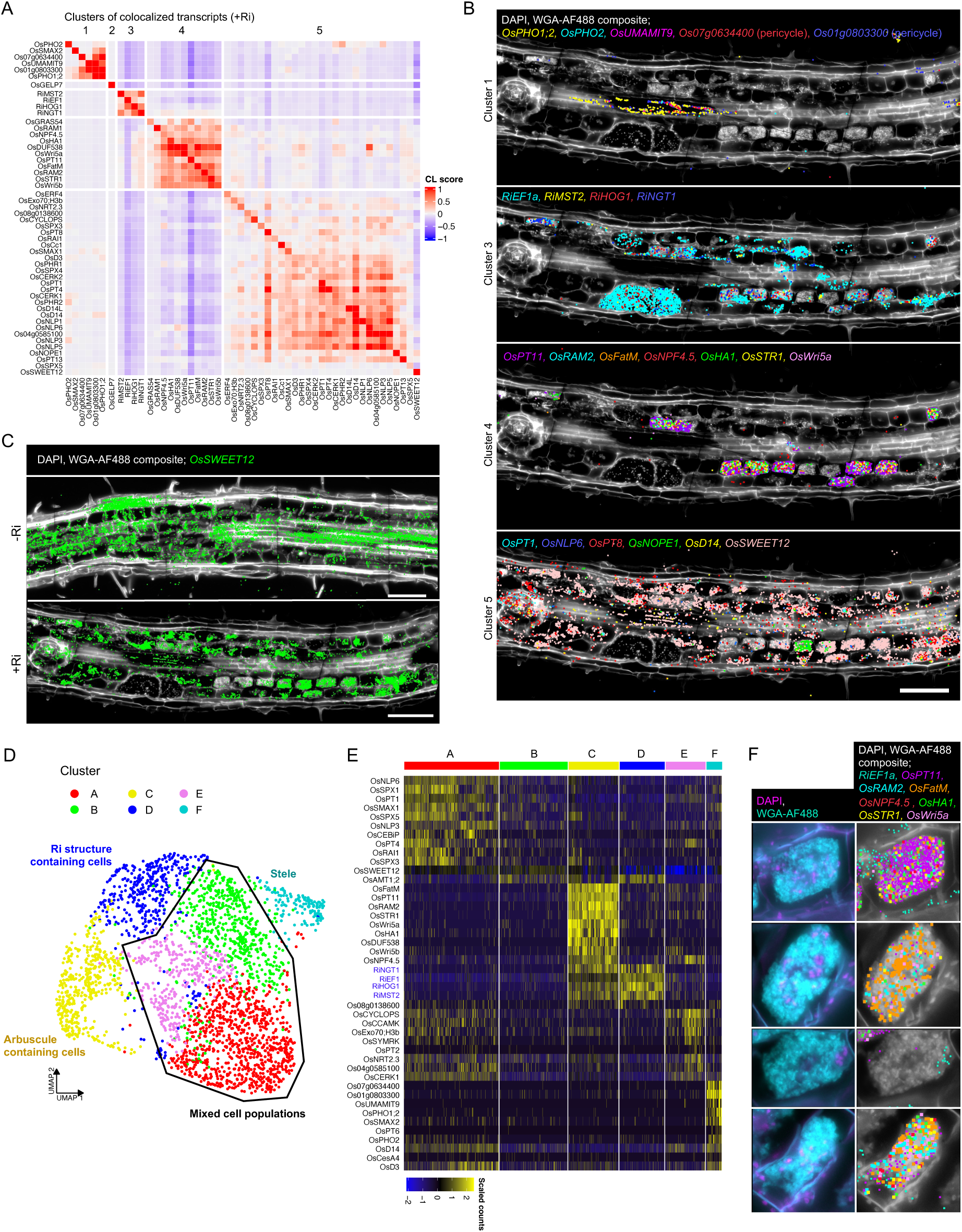
Co-localisation and cell segmentation analyses for Molecular Cartography mycorrhizal samples reveal distinct clusters of co-localised transcripts and cell populations. (A) Colocalization heatmap of transcript colocalization scores for +Ri sections. Genes with >0.2 self-co-localisation scores were used for analysis and subjected to hierarchical clustering. Distinct clusters are indicated by thicker boundaries between genes. Colour scale represents the transcript colocalization (CL) score. (B) Image-transcript overlays highlighting the spatial expression of key marker genes in each transcript colocalization cluster. (C) Image-transcript overlays highlighting spatial repatterning of *OsSWEET12* in colonized tissues. (D) UMAP projection of segmented cells, colour indicates the cluster (A to F) assigned to each cell by *Seurat* single-cell analysis. (E) Heatmap of normalised scaled transcript counts for each cell (columns) for the top 10 marker genes (rows) in each cell cluster (column sections), and their relative expression level compared to other clusters. (F) Image-transcript overlays demonstrating arbuscule individuality within a single root section; whereby phenotypically similar arbuscules exhibit distinct patterns of gene expression. DAPI/WGA-AF488 composite in white, different colours correspond to independent transcripts, each spot corresponds to one detected transcript. Scale bar, 100 μm.

Cluster 1 comprised vascular-associated targets, including *Os01g0803300*, *Os07g0634400*, xylem-loading phosphate transporter *OsPHO1;2*, and amino-acid transporter *OsUMAMIT9* (figure 3A, B). While exodermal markers clustered together in the mock condition, their reduced abundance in colonized roots left *OsGELP7* as the sole representative in cluster 2, possibly reflecting symbiosis-induced reprogramming of outer root layers. Cluster 3 contained fungal genes which formed a distinct group with no strong associations to rice targets. Cluster 4 was enriched for arbuscule-specific genes, including *OsSTR1*, *OsWRI5a/b*, and *OsDUF538*. Strong colocalization between *OsSTR1* and *OsWRI5b* reinforced their known roles in lipid transfer, while *OsDUF538* clustered tightly with *OsWRI5a*, suggesting a previously unrecognized role in arbuscule-associated lipid metabolism or signalling (figure 3A). Cluster 5 represented the largest gene group, containing AM and phosphate signalling genes (*OsSPXs*, *OsCERK1/2*, *OsD14*, *OsPHR1/2*), the phosphate regulator *OsPHO2*, and sugar transporter *OsSWEET12*. Although *OsSWEET12* transcript levels did not differ significantly between conditions (figure S2), spatial mapping showed redistribution in colonized roots, where expression shifted to arbuscule-containing cortical cells (figure 3C), consistent with localized sugar allocation to the fungus.

While transcript colocalization identified groups of genes with shared spatial patterns across tissues, this approach does not directly link expression profiles to individual cells. To resolve how these patterns map onto discrete cellular states and AM developmental stages, we next performed single-cell segmentation of colonized roots (figure S9A), generating an expression matrix of 3,464 cells. Vesicles were treated as single ‘cells’ due to their intercellular position. Six cell clusters were identified, which could be mapped back to their spatial location to identify the cell-type and AM developmental stage, facilitating annotation (figure 3D, 3E, S9B). Cluster C cells were the most consistent in both transcript abundance and AM developmental stage, with clear expression of arbuscule-localized plant genes as well as fungal transcripts, and with the majority of cells containing arbuscules, reflecting an intimate symbiotic state. Cluster D encompassed cells containing fungal structures (intraradical hyphae, vesicles) and showed high expression of fungal genes alongside plant *OsAMT1;2*, suggesting close coordination of ammonium transport (figure S10). Cluster F grouped mostly stele-associated cells, aligning with transcript colocalization cluster 1. However, clusters A, B and E cells were highly variable in terms of transcript abundance and AM development. Cluster A was enriched for symbiosis-related signalling genes (*OsSMAX1, OsNLP6, OsSPXs*), Cluster B cells did not express any distinct marker gene, and Cluster E cells displayed reduced *OsSWEET12* but elevated *OsCYCLOPS* and *OsNPF4.5*. More surprisingly, these clusters contained cells at different stages of colonization, ranging from uncolonized cortical cells, cells with developing/collapsing arbuscules, as well as uncolonized cells in the stele.

Therefore, while single-cell analysis of spatial transcriptomics data enabled the classification of distinct cell populations, it was not sufficient to accurately predict the AM developmental stage of individual cells. This may be explained by the inherently dynamic nature of AM symbiosis itself. Indeed, morphologically indistinguishable arbuscules within the same section displayed highly divergent expression signatures (figure 3B, 3F). This finding suggests that arbuscules are not uniform structures but display substantial individuality, with functional states shaped by local cellular context such as nutrient status, signalling flux, and host regulatory inputs. Together, these results show that AM colonization profoundly alters the spatial organization of transcripts across root cell types, redefining canonical markers and modulating nutrient and signalling pathways in a localized and heterogeneous fashion.

### 3.3 AM-inducible TRAP-seq for targeted investigation of translational responses throughout AM symbiosis

To complement our spatial transcriptomics dataset and address the challenge of capturing dynamic gene regulation across continuously shifting AM developmental stages, we applied stage-specific, AM-inducible TRAP-seq. TRAP-seq captures transcripts of distinct cell types or states that are associated with ribosomes containing an epitope-tagged Ribosomal Protein L18/u18 produced from a chimeric gene driven by a cell-type or conditional promoter (Mustroph et al., 2009b). We used this approach to monitor translating mRNAs during AM development in an unbiased manner. By isolating ribosome-bound transcripts, TRAP-seq provides a more direct proxy of active translation than bulk RNA-seq (Lee & Bailey-Serres, 2019). Three rice promoters known to be activated at different stages of AM colonization were used to construct TRAP lines (figure 4A). The *Arbuscular Mycorrhizal specific marker 1* (*OsAM1*) encodes a putative type III peroxidase PRX53, and is highly expressed during mycorrhizal colonisation (Güimil et al., 2005). *OsAM1* transcripts accumulate in cells containing small arbuscules and in cells flanking intercellularly growing hyphae (Gutjahr et al., 2008), and thus the *OsAM1* promoter is already active during early colonization, and remains active as new infection events arise during the asynchronous progression of symbiosis. The well-characterised *OsPT11* encodes an AM-specific phosphate uptake transporter which is expressed in the PAM surrounding fine-branched arbuscules, hence representing a “mid” colonization reporter marking the early-mature arbuscule development stages (Kobae & Hata, 2010; Paszkowski et al., 2002). Finally, the *Arbuscular Receptor-like Kinase 1* (*OsARK1*) encodes a receptor-like kinase (RLK) which regulates AM fungal fitness and plays a crucial role in symbiotic maintenance during post-arbuscule development. Translational reporter lines demonstrate that OsARK1 is localised to the PAM surrounding fine branches of mature and collapsing arbuscules, therefore representing a mid-late colonization reporter (Roth et al., 2018).

**Figure 4.**
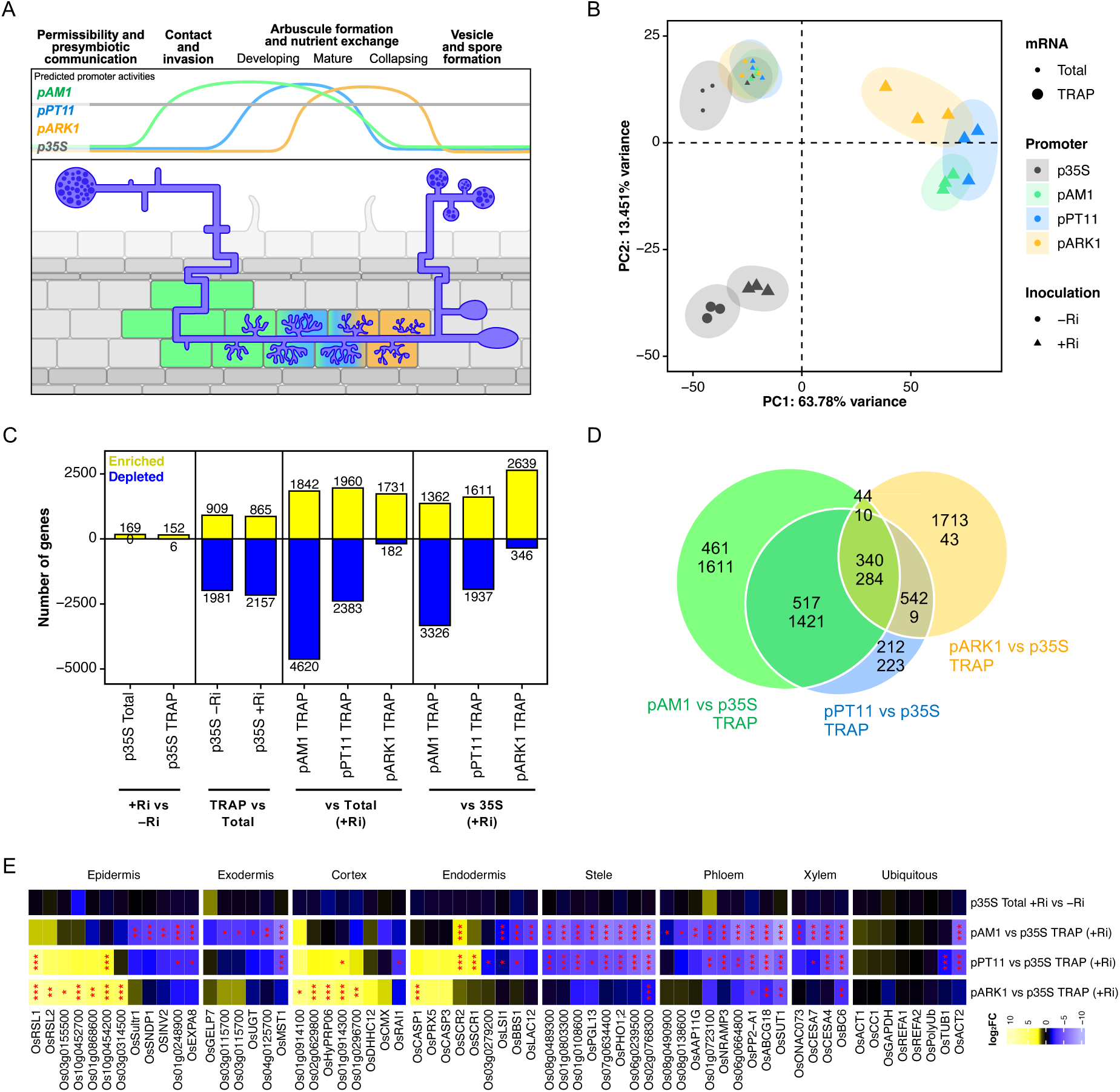
Stage-specific AM-inducible TRAP-seq shows dynamic translational reprogramming across AM development. (A) Schematic representation of predicted activities through root cell layers and AM development of the promoters used for AM-inducible TRAP. (B) Principal Component Analysis (PCA) visualization for all samples in the dataset, using the top 1000 variable features from normalised variance-stabilised-transformed (VST) gene counts. PC1 and PC2 displayed as explaining a combined 77.2% of the variance. Colour based on genotype, that is, promoter driving TRAP construct. Size based on total vs TRAP mRNA. Shape based on inoculation. Best fit ellipses for each treatment group were created using the Khachiyan algorithm from the *ggforce* package in R (Pedersen, 2025). (C) Number of differentially captured genes in all relevant comparisons between conditions. Positive yellow bars for up-regulated genes, negative blue bars for down-regulated genes. (D) Venn diagram showing overlaps between DEGs for each AM inducible promoter vs *p35S* TRAP samples, first number corresponds to up-regulated genes, second number to down-regulated genes. (E) Heatmap of log_2_ fold-changes (log_2_FC) for a selection of comparisons of interest, for a selection of cell-type specific marker genes (full comparison list and complete set of marker genes on figure S12). Significance levels as determined by DESeq2 shown with asterisks (* for p-value <0.05, ** for < 0.01, *** for < 0.001). Genes were subjected to hierarchical clustering within each group.

The three AM-inducible TRAP lines (*pAM1:TRAP, pPT11:TRAP, pARK1:TRAP)* were investigated alongside a constitutive TRAP line generated using the *CaMV35S* promoter (*p35S:TRAP*). To ensure uniform and extensive colonization, a nurse inoculation system was employed. For each line, both the TRAP fraction (ribosome-bound mRNAs) and the total mRNA pool were sequenced. Comparison of total and TRAP profiles from the constitutive *p35S:TRAP* line, grown under mycorrhizal conditions, established a baseline for differences between total and ribosome-associated mRNA populations in colonized roots. Applying the same analysis to the AM-inducible TRAP lines revealed additional changes reflecting both translational regulation and the enrichment of specific cell types targeted by each promoter. Finally, by comparing the TRAP profiles of each AM-inducible line with that of the constitutive *p35S:TRAP* line, we identified translational signatures specifically associated with distinct stages of AM development.

Principal component analysis (PCA) revealed clear separation of samples according to mycorrhizal status (+/–Ri), mRNA source (total or TRAP), and promoter used for TRAP lines (*pAM1, pPT11, pARK1, p35S*) (figure 4B). Along PC2 (13.5% variance), total versus TRAP samples of the constitutive *p35S:TRAP* line separated, reflecting differences between the transcriptome and translatome in whole tissue. PC1 (63.8% variance) primarily distinguished inoculated from non-inoculated samples, across both total and TRAP mRNA in the *p35S:TRAP* line, as well as between promoters driving TRAP expression. This axis therefore captured transcriptional and translational alterations associated with AM symbiosis and its cell populations. These patterns demonstrate that TRAP was effective in both constitutive and AM-inducible contexts, resolving promoter-specific translatomes distinct from the bulk transcriptome. Consistent with this, normalised gene counts showed strong enrichment of *OsAM1*, *OsPT11* and *OsARK1* in the TRAP fractions of their respective lines, an order of magnitude greater than in the *p35S:TRAP* line, confirming targeted enrichment (figure S11A).

Differential abundance analysis conducted on the total mRNA profiles of *p35S:TRAP* identified 169 upregulated genes in response to AM colonization (+Ri vs −Ri) (figure 4C) consistent with the general predominance of upregulated over downregulated genes in mycorrhizal systems (Fiorilli et al., 2013; Rich et al., 2017; Sgroi et al., 2024; Vangelisti et al., 2018). Of these, all but 15 overlapped with published bulk RNA-seq datasets (figure S11B). The relatively modest number of DEGs suggests that sampling captured an early stage of colonization, when key transcriptional programs initiating symbiosis are activated, while still encompassing cells at more advanced developmental stages. The same comparison performed on TRAP mRNA revealed 152 upregulated and 6 downregulated genes, with substantial overlap with the total mRNA set (109 shared) but also fraction-specific responses; for example, *OsNOPE1* was induced only in total mRNA, whereas *OsPT13* was specific to the TRAP fraction (figure S11B). Finally, direct comparison of TRAP and total mRNA pools within each colonization state revealed extensive differences (2,890 and 3,022 DEGs in –Ri and +Ri roots, respectively), two-thirds of which were independent of colonisation. The remaining third included translational shifts in strigolactone/karrikin signalling genes, such as *OsD53* and *OsSMAX1* (altered only +Ri), or *OsD53L* and *OsSMXL2* (altered only −Ri) (figure S11C). Collectively, these results show that AM symbiosis affects both transcriptional and translational regulation, largely through parallel expression changes but with distinct cases of translational control, consistent with observations in other biotic interactions (Traubenik et al., 2020; Traubenik et al., 2021).

We next compared the TRAP mRNA profiles of each AM-inducible line with (i) its corresponding total mRNA profile and (ii) the TRAP profile of the constitutive *p35S:TRAP* line. Both comparisons revealed significant enrichment of numerous genes, with comparable overall magnitudes of change, though enrichment tended to be slightly higher relative to the total mRNA profiles, except in the *pARK1* line. (figure 4C). This is consistent with both comparisons capturing transcripts enriched or depleted in the specific cell populations targeted, while the contrast with total mRNA additionally reflects translatome versus transcriptome differences. Importantly, the number of genes differentially captured in AM-inducible TRAP samples relative to *p35S* was an order of magnitude greater than in the +Ri vs −Ri comparison, indicating a strong enrichment of colonised cell populations. This is further supported by bulk mycorrhizal transcriptional response being largely recapitulated in the *pAM1* versus *p35S* TRAP comparison, with 147 out of 169 genes recovered (figure S11D). Moreover, when comparing with bulk RNA-seq data at a later co-cultivation time-point of six weeks, with higher colonisation levels (Das et al., 2022), further 529 genes transcriptionally activated by AM colonisation, not captured in our early bulk dataset, were significantly enriched in the *pAM1:TRAP* samples (figure S11D). This again supports specific capture of colonised cell populations, as the TRAP profile recapitulates the bulk transcriptomic response of roots with higher percentage of colonisation. Notably, beyond this established AM transcriptional response, more than half of the genes enriched for *pAM1:TRAP* samples, as well as a substantial down-regulation of genes absent from the total mRNA comparison, were uniquely captured by AM-inducible TRAP, revealing novel signatures associated with colonised root cells (figure S11D).

Comparing between AM-inducible lines, the number of enriched/depleted genes were in line with the expected timeframes for promoter activity (*pAM1* having the broadest range and *pARK1* the narrowest). *pAM1:TRAP* exhibited the greatest number of enriched transcripts (1588 up/4041 down), followed by *pPT11:TRAP* (1652 up/1957 down) and *pARK1:TRAP* (1402 up/106 down), consistent with these lines allowing to capture distinct cell-states or populations through AM development (figure 4C). Overlaps between these gene lists also followed the expected developmental trajectories of these cell populations. *pAM1:TRAP* and *pPT11:TRAP* shared the largest proportion (34.7%), while *pAM1:TRAP* and *pARK1:TRAP* shared the smallest (9.7%). *pPT11:TRAP* was mostly recapitulated by the other two lines, while *pAM1* and *pARK1* had substantial unique signatures. For *pAM1*, the majority of these were depleted, conversely for *pARK1* the majority were enriched, thus providing a unique and much-needed resource to investigate transcriptional reprogramming during early symbiotic interaction and arbuscule collapse (figure 4D).

To further validate the capture of distinct cell-populations by AM-inducible TRAP, we examined the expression patterns of a set of known cell-identity markers identified from the literature, including single-cell and spatial transcriptomic datasets in rice (Liu et al., 2021; Wang et al., 2021; Zhang et al., 2021; Zhu et al., 2025). The vast majority of cell-type specific genes were differentially regulated in the AM-inducible TRAP profiles, in contrast to ubiquitous controls such as *OsCc1* and to the total mRNA +Ri vs −Ri comparison (figure 4E, S12). Most markers were downregulated in *pAM1* and *pPT11* TRAP relative to total mRNA or *p35S*:TRAP, particularly those associated with vascular cell-types. This is consistent with TRAP capturing colonised cortical cells, where stele-specific transcripts being underrepresented. Interestingly, this trend is absent in *pARK1* TRAP comparisons. Instead, several cortical, exodermal, epidermal and endodermal markers appeared enriched both in *pARK1* and *pPT11*, but depleted in *pAM1*. These dynamic profiles cannot be explained by capture of cortical or epidermal cell types alone. Together with results from Molecular Cartography (figure S13), this suggests dynamic repatterning of cell-identity markers across AM development, with initial downregulation during early colonisation and arbuscule formation, followed by recovery during arbuscule collapse. Notably, many cell-type markers were also differentially represented in *p35S* TRAP and total mRNA pools, suggesting either translational regulation of cell-identity markers, or incomplete capture of all cell-types by *p35S:TRAP.* Finally, even within ubiquitous control genes, cytoskeletal components such as actin (*OsACT2*) and tubulin (*OsTUB1)* were significantly reduced in AM-inducible TRAP profiles, especially *pPT11*, consistent with the extensive cytological remodelling associated with arbuscule development (figure 4E, S12).

### 3.4 Temporal dynamics of gene activity across AM development

To identify biological processes dynamically regulated during symbiosis, we performed Gene Ontology (GO) term enrichment analysis on differentially captured genes from each AM-inducible *TRAP* line compared to *p35S*. This revealed stage-specific functional programs consistent with promoter activity. In *pAM1:TRAP* and *pPT11:TRAP*, enriched genes corresponded to AM-related processes such as “response to symbiotic fungus” and “lipid metabolism”, while *pARK1:TRAP* instead highlighted “cell wall organisation,” “pectin catabolism,” and “hydrogen peroxide catabolism” (figure 5A). This differential enrichment for *pARK1:TRAP* suggests substantial functional alterations during late stages of arbuscule development. Depleted genes in *pAM1:TRAP* and *pPT11:TRAP* included components of the strigolactone signalling pathway as well as biosynthetic enzymes, reflecting a biological shift in strigolactone biosynthesis and signalling through AM development (figure S14). Also decreased were transcripts of genes linked to stress and immunity, such as responses to jasmonic acid (JA) and oxidative stress. Conversely, stress-related terms (e.g. “hydrogen peroxide catabolism”, “response to other organism”) were enriched among *pARK1:TRAP* upregulated genes. Comparing depleted genes with those induced by pathogenic *Magnaporthe oryzae* infection (Yang et al., 2021) showed broad overlap for *pAM1:TRAP* and *pPT11:TRAP* but limited overlap for *pARK1:TRAP* (figure 5B). These included classical defence genes (encoding PR proteins, WRKYs, chitinases) and associated pathways (JA and ethylene signalling, flavonoid biosynthesis, redox regulation), supporting the idea that AM development fine-tunes immunity; downregulated during colonisation and maintenance, but reactivated during arbuscule collapse (figure S15A). Cell wall-related terms were similarly highly enriched in a stage-dependent manner. *pAM1:TRAP* and *pPT11:TRAP* showed depletion of cellulose synthases and expansins, while the latter were enriched in *pARK1:TRAP*, consistent with the requirement for wall loosening during arbuscule formation and remodelling during degeneration (figures 5A, 5C, S15B).

**Figure 5.**
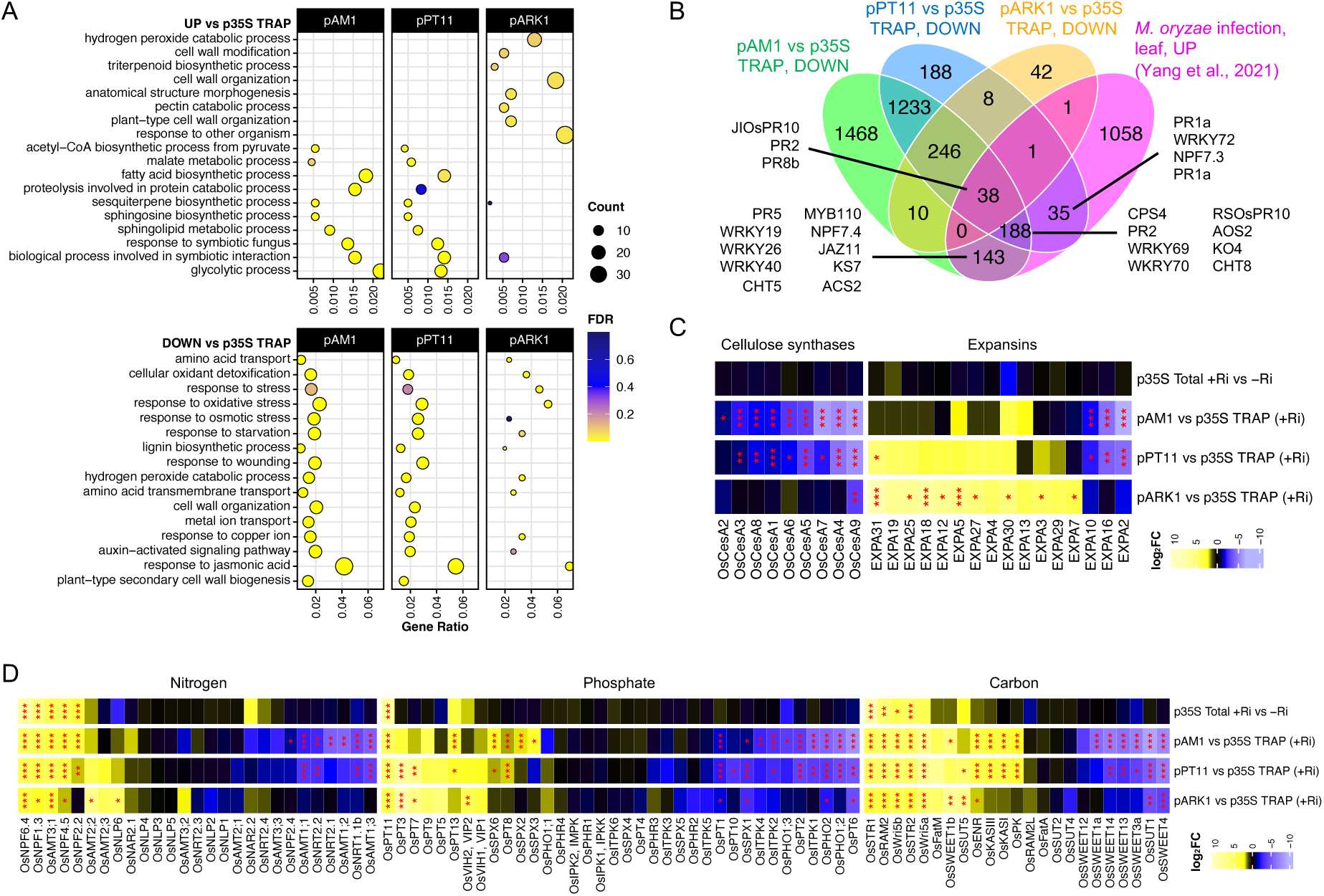
Dynamic regulation of biological processes, including defence, cell-wall biosynthesis and nutrient signalling, at specific stages of AM development. (A) GO term enrichment plot for up and downregulated genes in the translatomes obtained by use of three AM-inducible TRAP lines compared to the *p35S:TRAP* line. Top 10 GO terms by gene ratio (proportion of genes in dataset out of all known genes in the GO term) for each set are shown; colour scale indicates significance as false discovery rate (FDR); dot size proportional to the number of genes in each set included in the GO term. (B) Venn diagram showing overlaps between down-regulated genes for *pAM1*, *pPT11* and *pARK1* vs *p35S*:TRAP samples, compared to activated genes in *M. oryzae* infected leaves (Yang et al., 2021). (C) Heatmap of log_2_ fold-changes (log_2_FC) for a selection of comparisons of interest, for cellulose synthases and expansins (full comparison list on figure S15B). (D) Heatmap of log_2_FC for a selection of comparisons of interest (full comparison list on figure S16), for key genes related to phosphate, nitrogen and carbon transport, perception and signalling. Significance levels as determined by DESeq2 shown with asterisks (* for p-value <0.05, ** for < 0.01, *** for < 0.001). Genes were subjected to hierarchical clustering within each group.

We further investigated stage-specific profiles by identifying genes uniquely enriched or depleted under a particular promoter line. Among *pAM1:TRAP*-unique depleted genes (1611 in total), at least 138 encoded predicted transcription factors suitable as new regulatory candidates. In addition, at least 138 genes had predicted protein kinase activity, suggesting suppression of defence signalling cascades that might otherwise block colonisation. In *pARK1:TRAP*, uniquely enriched genes (1713 in total) included 185 genes involved in response to stimulus including 19 peroxidases, and at least 60 genes with predicted hydrolase activity. Four ribosomal small subunit (RPS) and six ribosomal large subunit (RPL) genes were also identified. Included were two *RPL23s*, a gene family known to be highly expressed under various abiotic stresses (Moin et al., 2016; Saha et al., 2017) and implicated in arbuscule collapse in *Medicago* (Floss et al., 2017). We also identified at least 59 genes encoding proteins with predicted TF activity, including an orthologue of the master regulator and maize-domestication gene *Teosinte-branched 1* (*Tb1*), named *OsTb2* or *OsREP1* (Lyu et al., 2020). These represent candidate genes regulating the post-arbuscule development stage.

Nutrient transporters and signalling components also showed interesting novel patterns. Beyond the induction of *OsPT11* and *OsPT13*, TRAP-seq revealed complex reprogramming of non-symbiotic phosphate transporters and signalling regulators undetectable in total mRNA profiles: *OsPT8* and *OsSPX2/6* were enriched in AM-inducible lines, whereas *OsPT1/2/6* and *OsPHO1;2/2* were depleted. Activation of *OsPT3* and *OsPT7* in *pPT11:TRAP* and *pARK1:TRAP*, and of *OsPT8* in *pAM1:TRAP* and *pPT11:TRAP*, further highlighted developmental resolution of phosphate signalling (figures 5D, S16). Similarly, nitrogen and carbon transporters (*OsAMTs, NRTs, SWEETs, SUTs*) displayed complex patterns of activation or repression in an AM-stage-specific manner. Many of these AM-regulated transcripts were not differentially abundant in bulk RNA-seq, particularly for genes repressed in early (*pAM1*) and mature (*pPT11*) stages. Comparison with our spatial transcriptomics dataset confirmed most nutrient-related trends, including depletion of *OsSWEET1a, OsSWEET14, OsPT6, OsPHO1;2, OsPHO2* and *OsSPX1*, which again were not detected as down-regulated in bulk RNA-seq (figure S13). However, spatial transcriptomics also revealed changes in spatial distribution, such as *OsSWEET12*, that were not associated with alterations in abundance at the transcript or translated mRNA levels (figure 3C, 5D, S13). Together, these complementary approaches demonstrate that AM symbiosis involves fine-scale, dynamic modulation of immunity, cell-wall remodelling, hormone signalling, and nutrient transport, with TRAP-seq providing developmental and translational resolution not captured by bulk transcriptomics.

## 4. Discussion

By integrating spatial transcriptomics with AM-inducible TRAP-seq, our study provides a high-resolution, cell-type-specific view of mRNA abundance and dynamics during symbiosis, offering new insights into the spatial and temporal coordination that underlies AM development.

Our spatial analysis surprisingly revealed transcript abundance for *RiEF1a*, an essential fungal translational elongation factor, was higher in vesicles and hyphae than in arbuscules, despite the latter’s central role in nutrient exchange. This discrepancy may be partially explained by differences in nuclear distribution. Vesicles and hyphae harbour visibly higher densities of fungal nuclei, likely supporting ongoing metabolic activity and cellular growth. In contrast, arbuscules, particularly their fine terminal branches, are structurally constrained, which may physically limit nuclear entry (Bianciotto et al., 1995). Rather than reflecting a reduced need for transcription, lower nuclear density in arbuscules may necessitate transcript or protein movement from the trunk and connected, adjacent fungal hyphae. This model is supported by previous knowledge from fungal species such as *Ustilago maydis*, where transcripts and proteins are actively transported over long distances within multinucleate hyphal networks (Zarnack & Feldbrügge, 2007, 2010). Therefore, transcription may be spatially decoupled from function, with vesicles and hyphae serving as biosynthetic hubs that support the specialized functions of arbuscules.

Despite detecting fungal transcripts within individual host cells, we observed minimal spatial colocalization between plant and fungal gene expression, indicating a high degree of functional compartmentalization even at the single-cell level. Intriguingly, we identified *OsAMT1;2*, a high-affinity ammonium transporter, in a discrete plant cell cluster expressing fungal genes but lacking classical arbuscule markers. Known to respond to local ammonium availability (X. Wu et al., 2022), *OsAMT1;2* mRNA in proximity to vesicles and hyphae suggests a role in sensing or scavenging nitrogen at these fungal interfaces, pointing to uncharacterized spatial dynamics in nitrogen acquisition.

Our analysis also uncovered reduced transcript abundance for most cell-identity markers in roots undergoing AM colonisation. This was further validated using TRAP-seq, where cell-identity markers showed significant alterations in their abundance in distinct cell-populations during AM symbiosis, particularly downregulated in early and middle stages, while reactivated during arbuscule collapse. These results both emphasize the need for systematic characterization of cell-type markers under abiotic and biotic stresses, in distinct root zones and cell types; as well as highlight that deep transcriptional and translated mRNA changes occur in AM-colonised tissue, even regarding cell-identity.

However, our spatial transcriptomic analysis failed to accurately predict AM developmental stages, particularly for the arbuscule-containing cell. This was most likely caused by a striking transcriptional heterogeneity among morphologically similar arbuscules. Even within the same tissue, arbuscules displayed marked differences in expression of key arbuscule localized genes such as *OsPT11*, *OsNPF4.5*, *OsSTR1*, and *OsRAM2*, suggesting that arbuscule function is “cell-autonomous”, likely fine-tuned in response to local physiological or developmental cues. These findings challenge the concept of a uniform “mature” arbuscule, and highlight the need to investigate arbuscule cellular heterogeneity, its regulatory causes, and functional consequences for nutrient transport.

AM-inducible TRAP-seq complemented spatial transcriptomics, providing developmental resolution by capturing ribosome-associated transcripts in colonized cell populations at distinct progressive stages of the interaction. Furthermore, while spatial transcriptomics revealed where genes are transcribed, TRAP-seq provided a complementary perspective by identifying which transcripts that had been recruited for translation. This approach uncovered extensive and dynamic translational reprogramming, particularly involving immunity, cell-wall and stress-related genes, which were consistently downregulated during early stages of colonization, and upregulated in later stages. Within this broader pattern, genes associated with JA signalling were strongly downregulated in early and mid colonization. Although JA is typically linked to defence responses, its contribution to AM symbiosis has been debated. For example, exogenous JA suppresses colonization in tomato (Herrera-Medina et al., 2008), whereas in rice, endogenous JA biosynthesis is dispensable for symbiosis, though elevated JA levels during pathogen challenge can inhibit fungal entry and development (Gutjahr, Siegler, et al., 2015). Our data help reconcile these apparently mixed reports: both JA and cell-wall–related pathways require dynamic regulation across AM development, initially suppressed to enable fungal entry and accommodation, then re-activated during arbuscule degradation.

Beyond immune modulation, our TRAP-seq data revealed dynamic regulation of genes involved in nutrient transport and signalling during arbuscule maintenance. As expected, symbiotic phosphate transporters such as *OsPT11* and *OsPT13* were enriched at the transcript and translated mRNA levels. However, AM-inducible TRAP-seq identified uncharacterised modulation of non-canonical phosphate transporters and signalling components, both induced (*OsPT3*, *OsPT7*, *OsPT8, OsSPX2, OsSPX6*) and repressed (*OsPT1*, *OsPT2*, *OsPT6; OsPHO1;2; OsPHO2*), in some cases dynamically through AM development, reflecting a stage-specific and nuanced control of phosphate uptake. Similar regulation, also undetectable in bulk RNA-seq, was observed for nitrogen and carbon transporters, including *OsAMTs*, *OsNRTs*, *OsSWEETs*, and *OsSUTs*. These findings highlight the pivotal role of local dynamic control in fine-tuning host nutrient management throughout the symbiotic timeline. By integrating TRAP-seq with spatial transcriptomics, we were able to contextualize the translatome changes with cell-specific mRNA patterns. While overall trends were consistent across methods, spatial transcriptomics uncovered localized gene expression nuances, exemplified by the striking spatial repatterning of *OsSWEET12* in colonized versus uncolonized regions.

During later stages of AM fungal development, we observed a coordinated upregulation of immunity and stress-related genes, indicative of arbuscule degeneration. This transition echoes findings in *Medicago truncatula*, where arbuscule collapse was associated with reactivation of defence pathways and cellular recycling processes (Floss et al., 2017). Importantly, TRAP-seq uncovered novel candidate genes potentially involved in late stages of arbuscule development and collapse. For instance, the large ribosomal subunit gene *OsRPL23* showed marked enrichment in late-stage colonized cells and shares homology with a Medicago gene previously implicated in arbuscule turnover (Floss et al., 2017). In addition, *OsREP1/OsTB2*, orthologue of maize-domestication gene *Tb1* encoding a TCP-family transcription factor, was one of the most highly enriched genes during the post-arbuscule development stage, suggesting a potential regulatory role in fungal withdrawal or reactivation of host developmental programs.

Together, our complementary approaches provide distinct but intersecting views of AM symbiosis. While the microbe programmed cortical cell translatome reveals actively translated genes within defined cell populations, spatial transcriptomics reveals expression landscapes within the broader tissue context. Integrating these perspectives reshapes our understanding of gene regulation during arbuscule development, function, and turnover, revealing a complex, multilayered regulatory network. Ultimately, our findings redefine key aspects of AM symbiosis by revealing transcriptional and translational heterogeneity across fungal structures and host cells, identifying immune reprogramming as a central feature of early accommodation, and uncovering finely tuned spatial regulation of nutrient transporters. Moving forward, combining these insights with proteomic analyses, live-cell imaging, and targeted genetic studies will be essential for mapping protein localization, validating candidate regulators, and decoding the molecular dialogue underpinning arbuscule development.

## 5. Materials and methods

### 5.1 Plant growth and inoculation

Seeds of *Oryza sativa cv. Japonica*, Nipponbare were surface-sterilized briefly in 70% (*v/v*) ethanol, then for 20 min in 3% (*v/v*) sodium hypochlorite. Imbibed seeds were germinated on 0.9% (*w/v*) bactoagar at 30°C for 7 days. Pre-germinated seedlings were transferred into pots (11 x 8 cm) containing sterile quartz sand in walk-in growth chambers at 12-h:12-h, light:dark cycle at 28:20°C and 60% relative humidity. Plants were inoculated with 900 spores of *R. irregularis* DAOM197198 (Mycorise® ASP, Premier Tech Biotechnologies, Rivière-du-loup, Canada). Plants were watered three times weekly for the first two weeks post-inoculation (wpi), thereafter fertilized twice a week with half Hoagland solution (25 µM Pi) and 0.01% (*w/v*) Sequestren Rapid (Syngenta), as previously described by (Gutjahr et al., 2008). Plants required for spatial transcriptomics experiments were harvested at 6 wpi.

For TRAP-seq experiments, 6 wpi wild-type plants were retained as “nurse plants” to enable rapid colonization of young test plants. Here, five pre-germinated seedlings were sown into pots, each already containing a colonized nurse plant. Test plants were harvested at 3 wpi.

### 5.2 Molecular Cartography™

#### Molecular Cartography probe design

Probes were designed using Resolve BioSciences’ proprietary design algorithm and gene annotations from the *Oryza sativa* ssp. *japonica* Nipponbare reference genome (Os-Nipponbare-Reference-IRGSP-1.0.58) (Kawahara et al., 2013). Searches were confined to the coding regions to identify potential off-target sites. Each target sequence underwent a single scan for all k-mers, favouring regions with rare k-mers as seeds for full probe design. A probe candidate was generated by extending a seed sequence until reaching a certain target stability. After these initial screens, probes were aligned with the background transcriptome, and probes with stable off-target hits were discarded. From the pool of accepted probes, the final set was composed by selecting the highest scoring probes.

#### Identification and Preparation of AM-Colonized Root Tissue

To facilitate the identification of AM-colonized root regions, we used a *pSCAMP:eGFP-SCAMP* rice reporter line expressing a GFP-tagged secretory carrier membrane protein, generated by (Kobae & Fujiwara, 2014). Roots were harvested at 6 wpi and screened for eGFP fluorescence using a Leica M205 FA stereomicroscope. GFP-positive root segments (0.3–0.5 cm in length) were immediately processed for spatial transcriptomics.

#### Tissue Preparation and Molecular Cartography

Root segments were fixed in paraformaldehyde (PFA) and embedded in paraffin according to Resolve Biosciences’ standard plant tissue preparation protocol, adapted from (Zöllner et al., 2021). Semi-thin sections (10 µm) were made using a Leica HistoCore Autocut R microtome and adhered to the Molecular Cartography slides. Sections underwent deparaffinization, permeabilization and refixation according to Resolve BioSciences’ user guide. Samples were mounted in SlowFade Diamond Antifade Mountant (Thermo Fisher Scientific) and shipped to Resolve Biosciences for Molecular Cartography analysis.

At Resolve BioSciences, sections were washed twice in 1×phosphate buffered saline (PBS) for 2 min, followed by 1 min washing in 50 % and 70 % ethanol at room temperature. Ethanol was removed by aspiration and DST1™ buffer was added, followed by tissue priming for 30 min at 37 °C and by a 48h hybridization using probes specific for the target genes. Samples were washed to remove excess probes and fluorescently tagged in a two-step colour development process. Fluorescent signals were removed after imaging in a decolourization step. Colourization, imaging, and decolourization were iterated for multiple cycles to generate a unique combinatorial code for each target gene. Samples were imaged by Resolve BioSciences using a Zeiss Celldiscoverer 7 with a 50×Plan Apochromat water immersion objective having a numerical aperture (NA) of 1.2 and a 0.5×magnification changer, resulting in a final magnification of 25× (Glenn et al., 2023). To visualize fungal structures, sections from experiment 1 were stained with WGA-AF633 (Invitrogen), although staining was suboptimal. In experiment 2, WGA-AF488 was used to improve fungal structure visibility. All sections were counterstained with DAPI to visualize both plant and fungal nuclei. Probes yielding fewer than 20 transcript counts per section across both −Ri and +Ri conditions were excluded from downstream analyses. Additionally, root sections with significant tissue damage or low overall transcript abundance were omitted from further analysis. Additionally, root sections with significant tissue damage or low overall transcript abundance were excluded from downstream analyses.

#### Image and Transcript Analysis

Image analyses, including generation of image-transcript overlays and manual cell-boundary segmentation was performed in Fiji under ImageJ software license (Schindelin et al., 2012). Transcript colocalization analysis was carried out using the Polylux plugin (Resolve Biosciences). Default settings were used for transcript colocalization analyses (xy-step size = 1, z-step size = 3.5, search radius = 50, and distance penalty = 4). To account for section-to-section variability, we calculated the median transcript colocalization score separately for all transcripts within mock and mycorrhizal samples. Single-cell clustering analysis was performed using the Seurat package in R. Cell cluster identities were then mapped back to their original spatial context in ImageJ using the Polylux plugin (Resolve Biosciences).

### 5.3 Translating Ribosome Affinity Purification (TRAP) and RNA-seq

*Promoter:TRAP* constructs were prepared with the TRAP destination vector *pH7WG-OsTRAP* which was engineered by modifying *p35S:HF-OsRPL18* (Reynoso et al., 2022). This vector contains a Gateway recombination site for promoter insertion upstream of a chimeric ribosomal protein fusion comprising a 6×His-FLAG-3×Gly tag, GFP, and OsRPL18/eL18 (*Os03g0341100*). Promoters listed were cloned into pENTR-D/TOPO (Invitrogen) and subsequently recombined into the TRAP vector via LR Clonase II enzyme mix (Invitrogen). All constructs were validated by Sanger sequencing. Resulting T-DNA plasmids were introduced into *Oryza sativa japonica* cv. Nipponbare embryogenic calli, derived from mature seed embryos, through *Agrobacterium tumefaciens-*mediated transformation. Callus for transformation of rice cultivar Nipponbare was generated by plating surface-sterilised mature seed, with embryo axes removed, on N6DT medium (3.95 g/L N6 basal salts, 30g/L sucrose, 300mg/L casein hydrolysate, 100mg/L myo-inositol, 2878mg/L proline, 0.5mg/L nicotinic acid, 0.5mg/L pyridoxine HCl, 1mg/L thiamine HCl, 37.3 mg/L Na_2_EDTA, 27.8 mg/L FeSO_4_, 2mg/L 2,4-D Na salt, 150mg/L Timentin, 4g/L Gelrite, pH5.8). Plates were sealed with Parafilm and cultured in the dark at 28C for 21 days after which time callus was cut into 2-4mm pieces, plated on fresh N6DT and cultured as before for a further 4 days. Preparation of Agrobacterium strain EHA105 containing the binary constructs and transformation of the rice callus pieces was carried out as previously described (Choi et al., 2020). T2 generation plants were used for TRAP experiments.

TRAP was performed as previously described (Mustroph et al., 2009a; Reynoso et al., 2015) with modifications outlined by Reynoso et al. (2019). From each sample, both total RNA and TRAP RNA (following ribosome purification) were extracted using the RNeasy Micro Kit (Qiagen). Libraries were prepared with the QuantSeq 3’ mRNA-seq kit (Lexogen) incorporating unique dual indices (UDIs) and unique molecular identifiers (UMIs) for three biological replicates per treatment. Sequencing was performed on an Illumina NovaSeq 6000 platform generating 150 bp single-end reads. Sequencing data was analysed by the Lexogen NGS Kangaroo Data Analysis Platform including quality control, trimming, UMI deduplication, STAR alignment to the *O. sativa* Nipponbare reference transcriptome (Os-Nipponbare-Reference-IRGSP-1.0.58) (Kawahara et al., 2013) and count quantification. To address low read counts in AM-inducible TRAP samples, an additional library was constructed and sequenced from the original RNA extractions, with resulting counts combined at the count level. Subsequent data analysis, including normalisation, filtering and differential gene expression, was carried out in R using the DESeq2 v1.40.2 package (Love et al., 2014), with a threshold of fold-change ≥ 1.5 or ≤ −1.5 and Benjamini-Hochberg false discovery rate (FDR) corrected *P*-value < 0.05. Gene Ontology (GO) enrichment analyses were conducted using the R package clusterProfiler v4.8.3 (Wu et al., 2021). The annotations of the genes including associated GO terms were collected from various sources: The Rice Annotation Project (RAP) (Kawahara et al., 2013) and EnsemblPlants (Martin et al., 2023) annotations for the Os-Nipponbare-Reference-IRGSP-1.0.58 reference genome and Oryzabase (Kurata & Yamazaki, 2006).

## 6. Data availability

Raw sequences, images and gene count matrices generated in this study are available in the NCBI GEO database under the accession GSE283774. R code employed to analyse the data and generate graphics is available in the following github repository: https://github.com/gabriel-ferreras/Spatial_rice_AMF_2025

Other data supporting the findings of this article are available in the supplemental information.

## Acknowledgements

We are grateful to Siobhan Brady, Neelima Sinha, Roger Deal, Alex Borowsky and Alex Canto-Pastor for helping shape the project, valuable discussions, and suggestions regarding data analysis. We would also like to thank colleagues at the Crop Science Centre, University of Cambridge for assistance with harvesting and constructive feedback, in particular Min-Yao Jhu for help with sectioning and spatial transcriptomics, Edwin Jarratt-Barnham, Liam German and Jen McGaley for assistance in gene list compilation, and Martina Orvosova for help in designing the graphical abstract. Research was supported by the National Science Foundation (NSF IOS-1856749 and IOS-211980, T.C., J.B.S. and U.P.), United States Department of Agriculture (2022-67012-36716, G.Z.A), Biotechnology and Biological Sciences Research Council (BB/P003176/1, E.W.; BB/P003419/1, U.P.), St John’s College (University of Cambridge) through the Benefactor’s Scholarship to G.F.G and Research Reimbursement Schemes to U.P., Cambridge Philosophical Society through Research Studentship to G.F.G.

