## Supplemental Information for "Spatiotemporal regulation of arbuscular mycorrhizal symbiosis at cellular resolution"

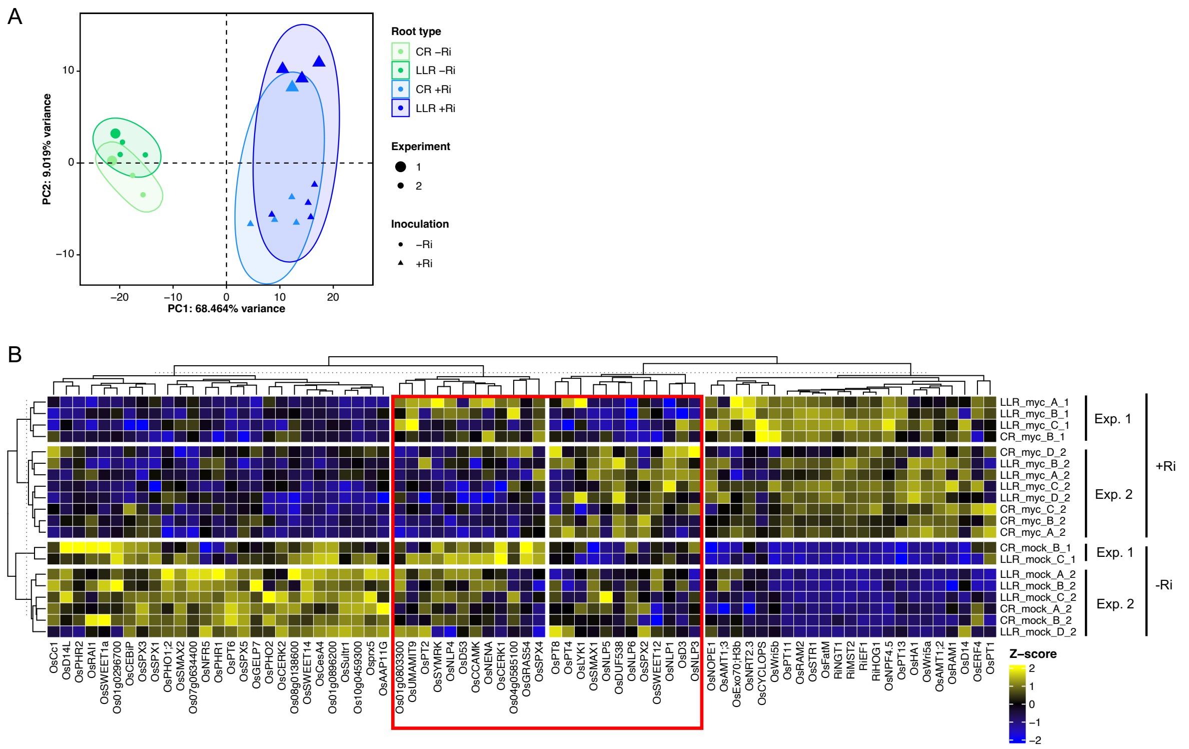


**Figure S1. Validation of Molecular Cartography results.** (A) Principal Components Analysis (PCA) for spatial transcriptomic sections, separated based on root type (colour, light for CR and dark for LLR), experiment (size) and inoculation (shape, circles for -Ri, triangles for +Ri). Variance-stabilised counts (VST counts) for transcript spot data for all filtered probes for each section were used to conduct PCA analysis. Only PC1 and PC2 are shown as they together account for more than 50% of the variance. Best fit ellipses for each sample type were created using the Khachiyan algorithm from the *ggforce* package in R (Pedersen, 2025). (B) Heatmap of VST row-centered counts for transcript spot data of each section. Genes and sections were subjected to hierarchical clustering followed by k-means partitioning to subdivide the set in four groups. CR, crown root; LLR, large lateral root; Ri, *Rhizophagus irregularis* inoculation.

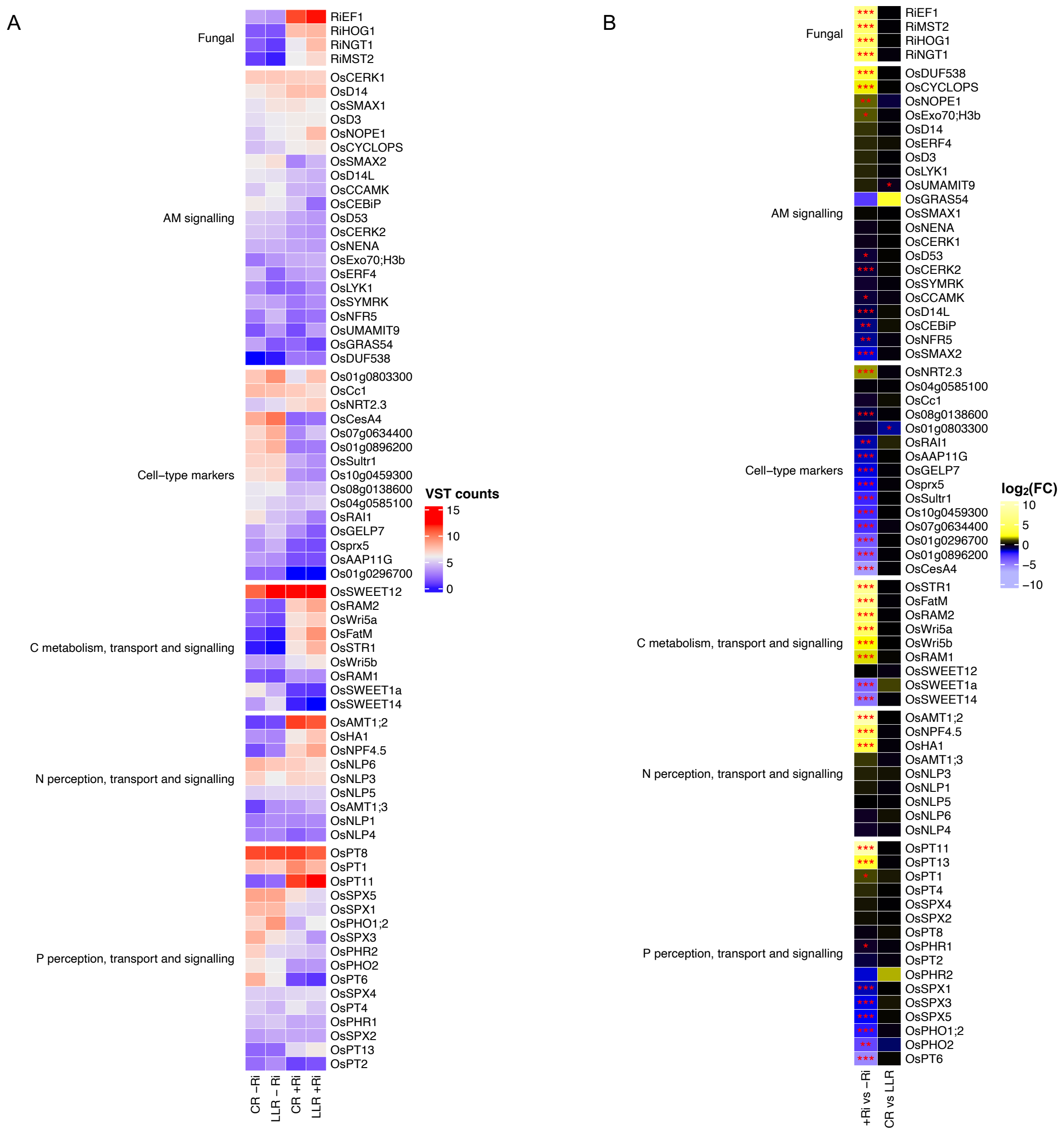


**Figure S2. Transcript quantification and differential gene expression analyses for Molecular Cartography results.** (A) Heatmap of VST counts for averaged transcript spot data for all sections of each condition (root type x inoculation). (B) Heatmap of $\log_2$ fold-changes ($\log_2$FC) for comparisons of transcript spot data between inoculation regimes and root types. Significance levels as determined by DESeq2 shown with asterisks (* for p-value <0.05, ** for < 0.01, *** for < 0.001). CR, crown root; LLR, large lateral root; Ri, *Rhizophagus irregularis* inoculation.

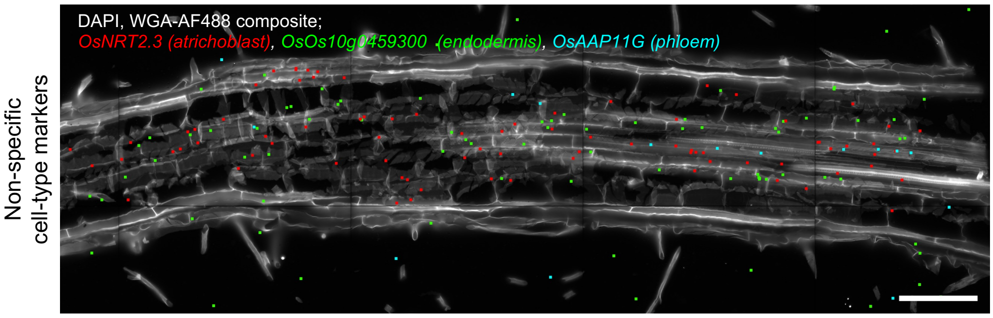


**Figure S3. Image-transcript overlay image of a non-inoculated (-Ri) section showing spatial expression of a selection of cell-type markers that were not specific** to the predicted cell-type in our samples. DAPI/WGA-AF488 composite in white, different colours correspond to independent transcripts, each spot corresponds to one detected transcript. Scale bar, 100 µm.

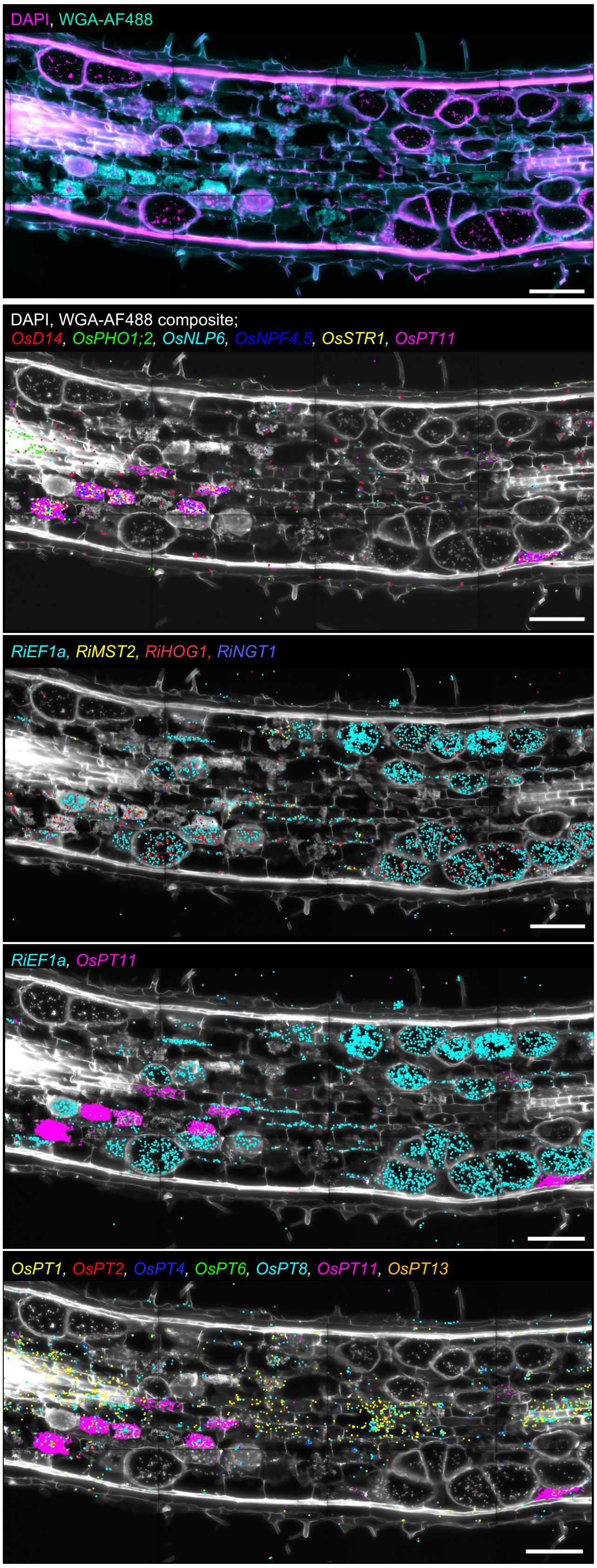


**Figure S4. Molecular Cartography results for an inoculated (+Ri) section.** First panel shows composite image of DAPI (magenta) and WGA-AF488 (cyan) for visualization of nuclei and cell boundaries. Rest of panels show image-transcript overlays for a selection of plant genes, fungal genes, one fungal and one plant gene, and plant phosphate transporters, in that order. DAPI/WGA-AF488 composite in white, different colours correspond to independent transcripts, each spot corresponds to one detected transcript. Scale bar, 100 µm.

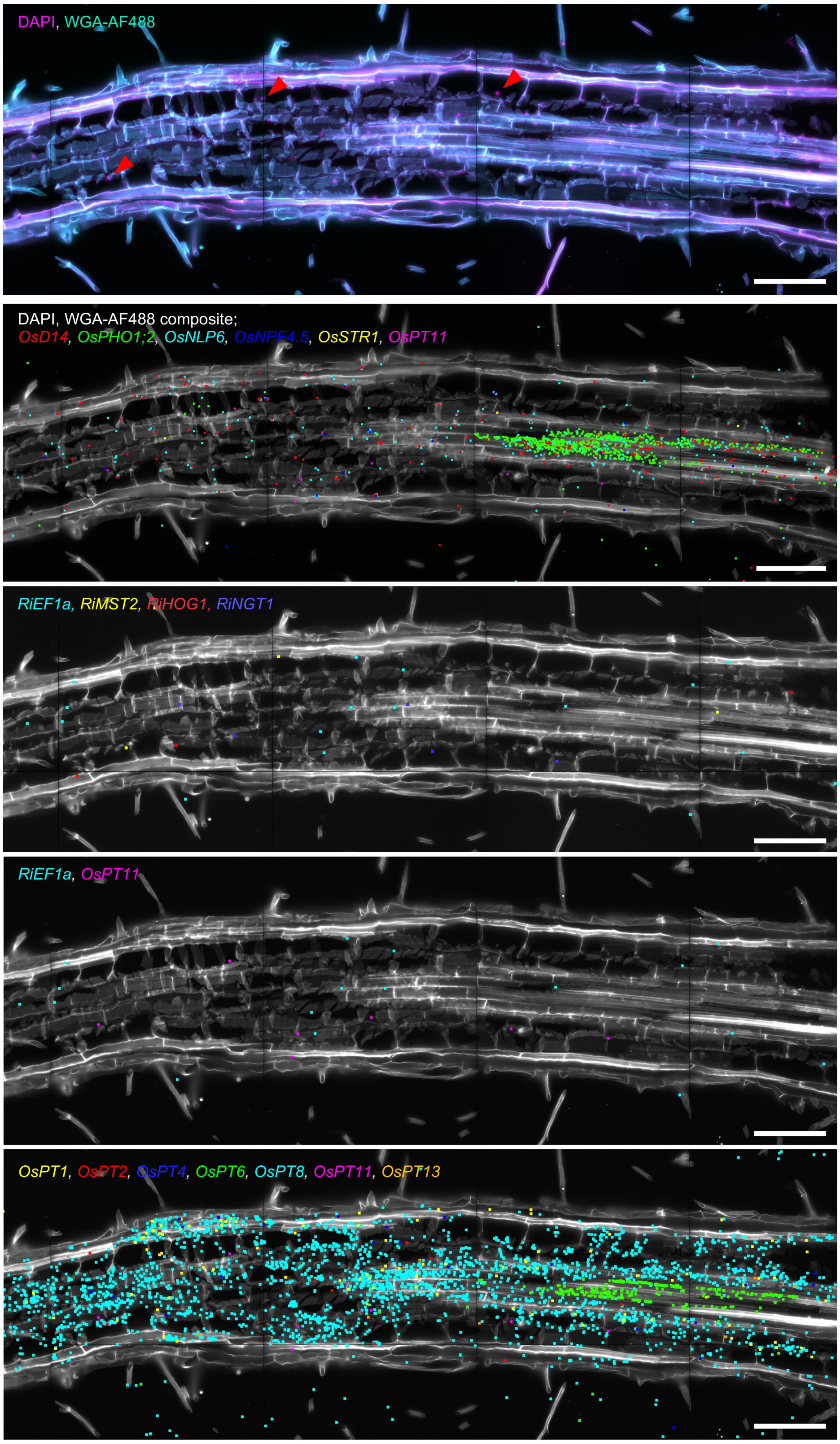


**Figure S5. Molecular Cartography results for an non-inoculated (-Ri) section.** First panel shows composite image of DAPI (magenta) and WGA-AF488 (cyan) for visualization of nuclei, cell boundaries and fungal structures. Rest of panels show image-transcript overlays for a selection of plant genes, fungal genes, one fungal and one plant gene, and plant phosphate transporters, in that order. DAPI/WGA-AF488 composite in white, different colours correspond to independent transcripts, each spot corresponds to one detected transcript. Scale bar, 100 μm.

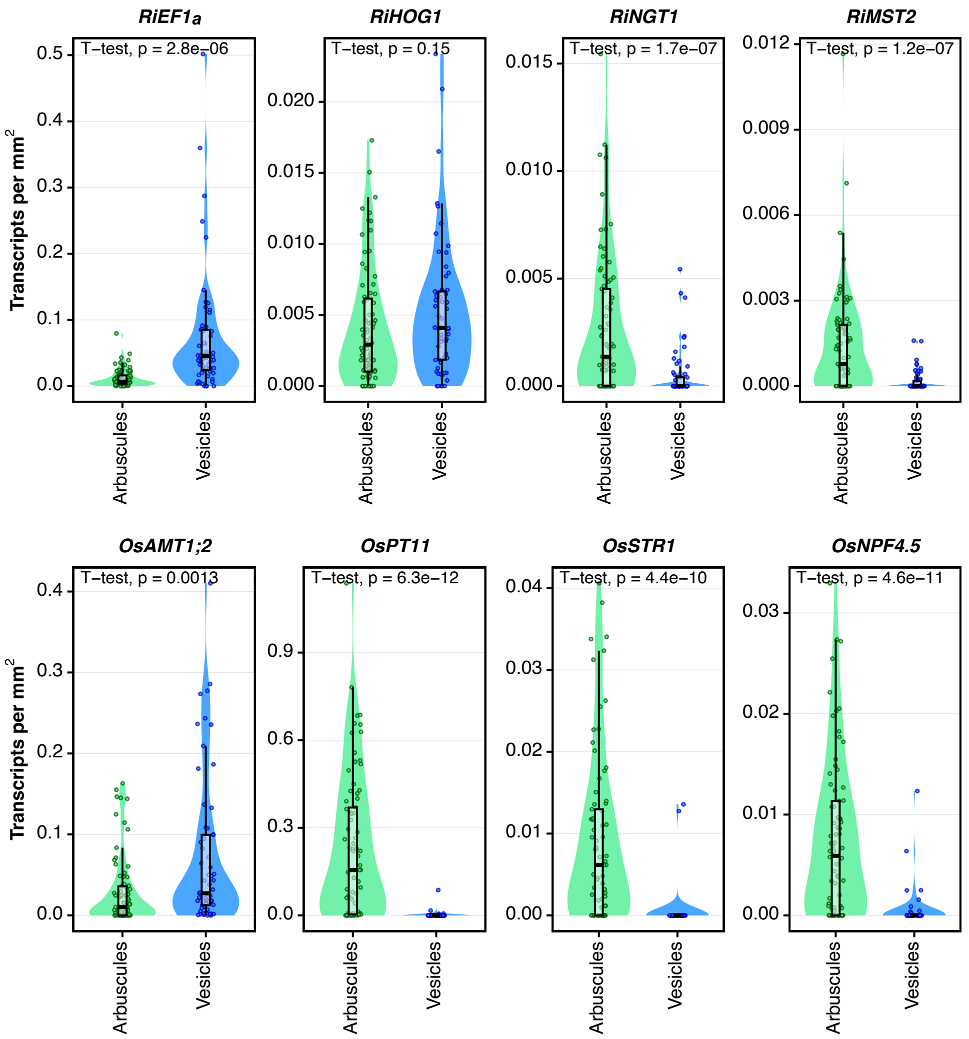


**Figure S6. Quantification of transcript abundance in distinct *R. irregularis* colonisation structures in Molecular Cartography +Ri sections**. Each graph corresponds to one transcript, name above. P-value of T-student test between structures indicated in each graph. Each spot corresponds to the transcript per $mm^2$ value for an individual structure (arbuscule or vesicle), n=96 for arbuscules, n=67 for vesicles.

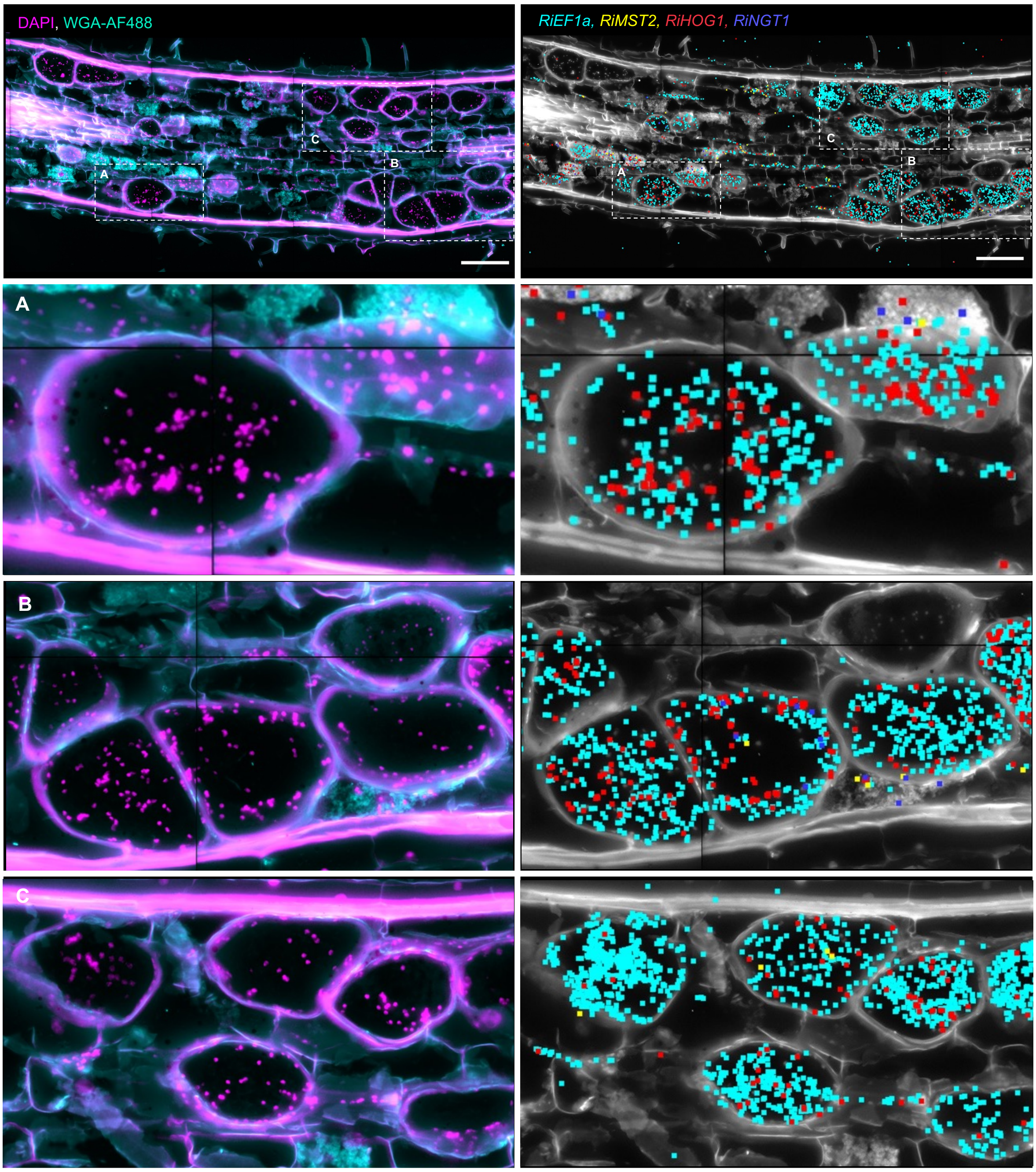


**Figure S7. Association between fungal nuclei and transcripts in vesicles of a +Ri section.** Left panels show composite image of DAPI (magenta) and WGA-AF488 (cyan) for visualization of nuclei, cell boundaries and fungal structures. Right panel shows image-transcript overlays for fungal genes; DAPI/WGA-AF488 composite in white, different colours correspond to independent transcripts, each spot corresponds to one detected transcript. First row for full section, following rows A, B and C for magnified vesicles, location indicated on first row. Scale bar, 100 µm.

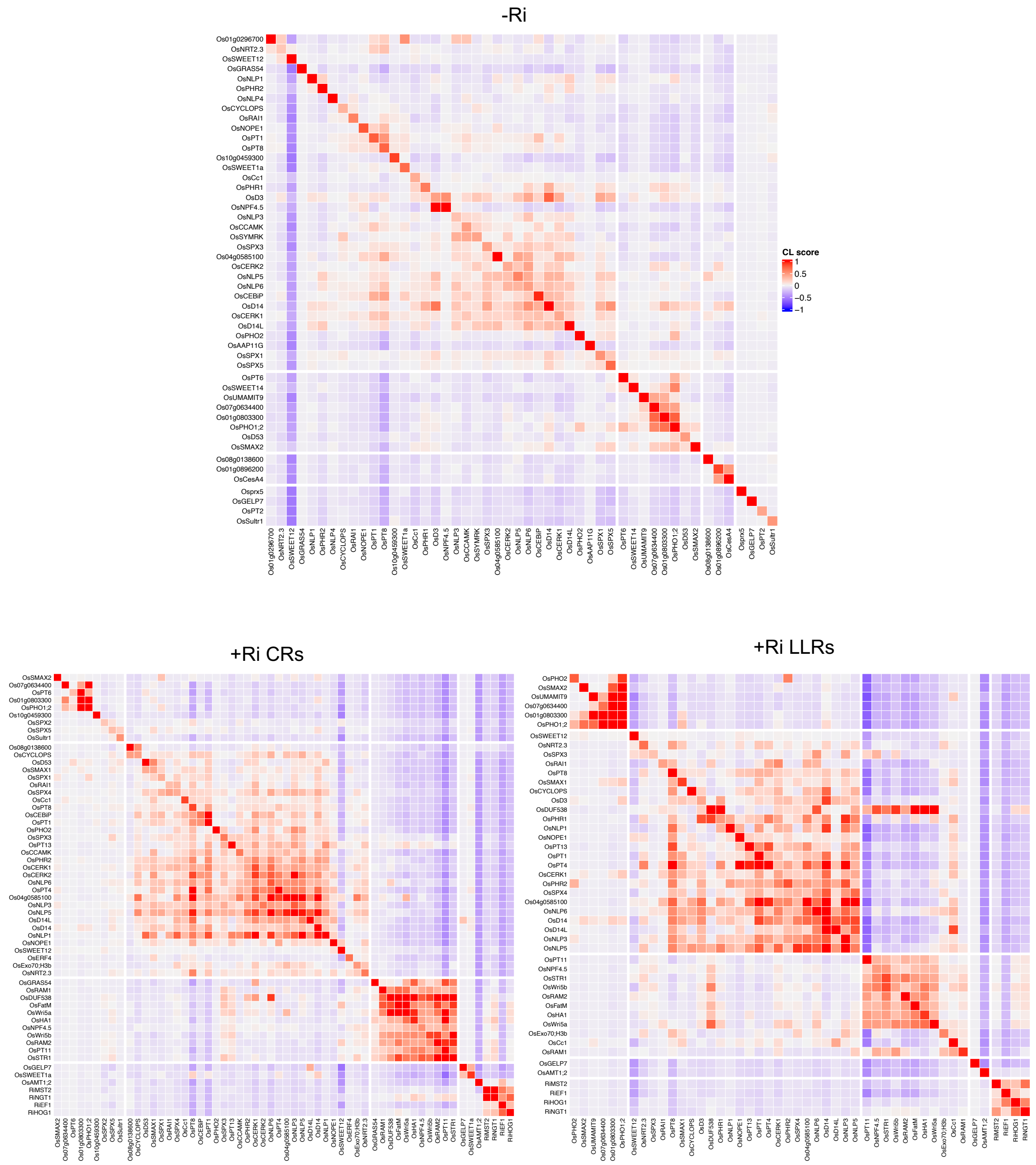


**Figure S8. Colocalization heatmap of transcript colocalization scores** for -Ri sections (A), +Ri crown root sections (B) and +Ri large lateral root sections (C). Genes with >0.2 self-co-localisation scores were used for analysis, subjected to hierarchical clustering, distinct clusters indicated by thicker boundaries between genes. Colour scale represents the transcript colocalization (CL) score.

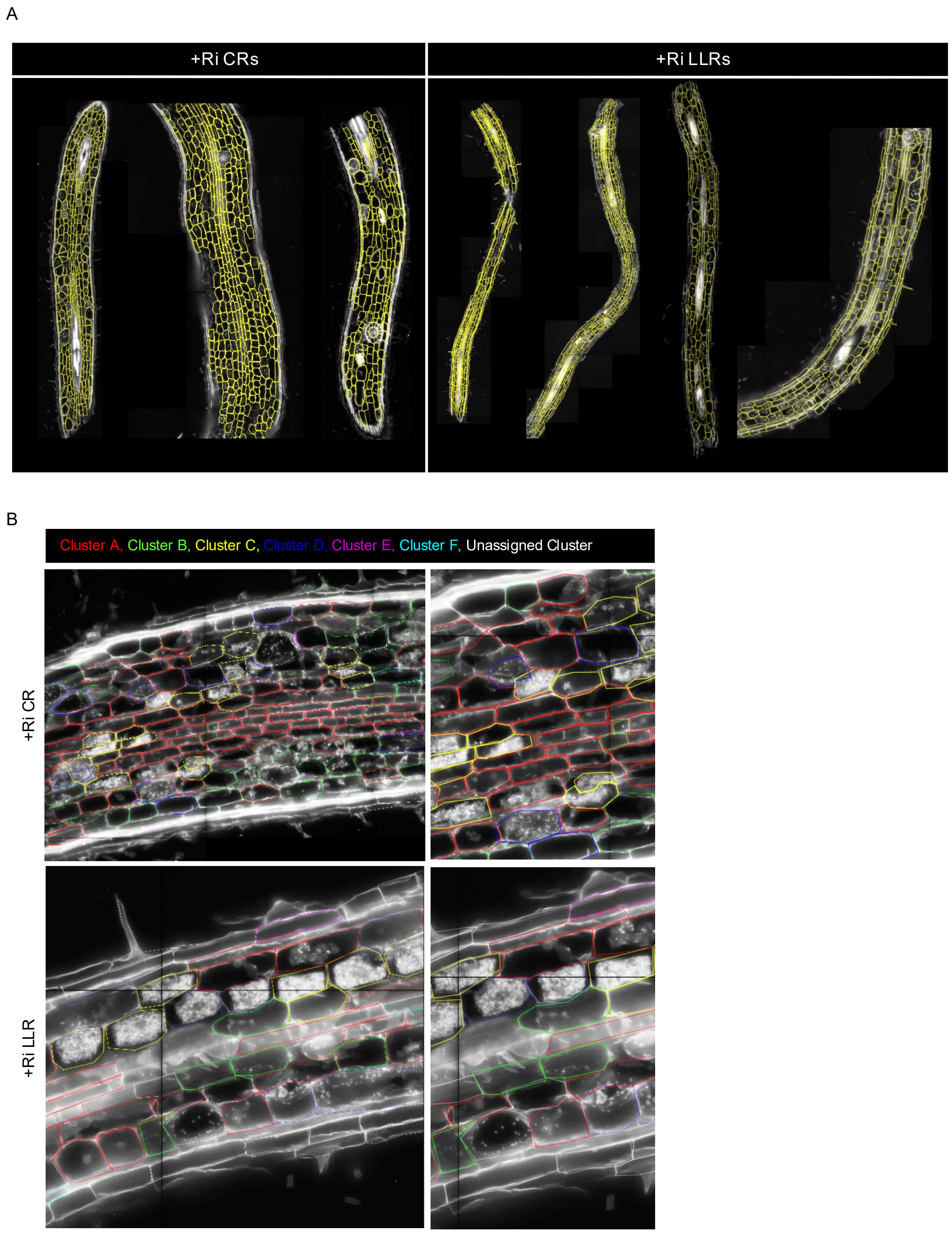


**Figure S9. Cell segmentation of Molecular Cartography sections and assignment of segmented cells to clusters.** (A) Cell-segmentation of Molecular Cartography +Ri sections, yellow lines indicate segmented cells, DAPI/WGA-AF488 composite in white. (B) Assignment of clusters to segmented cells based on Seurat single-cell analyses. DAPI/WGA-AF488 composite in white, segmented cells in coloured lines, distinct colours for separate clusters.

First row for a crown root section, second row for a large lateral root section. Right panels for zoomed-in close-ups.

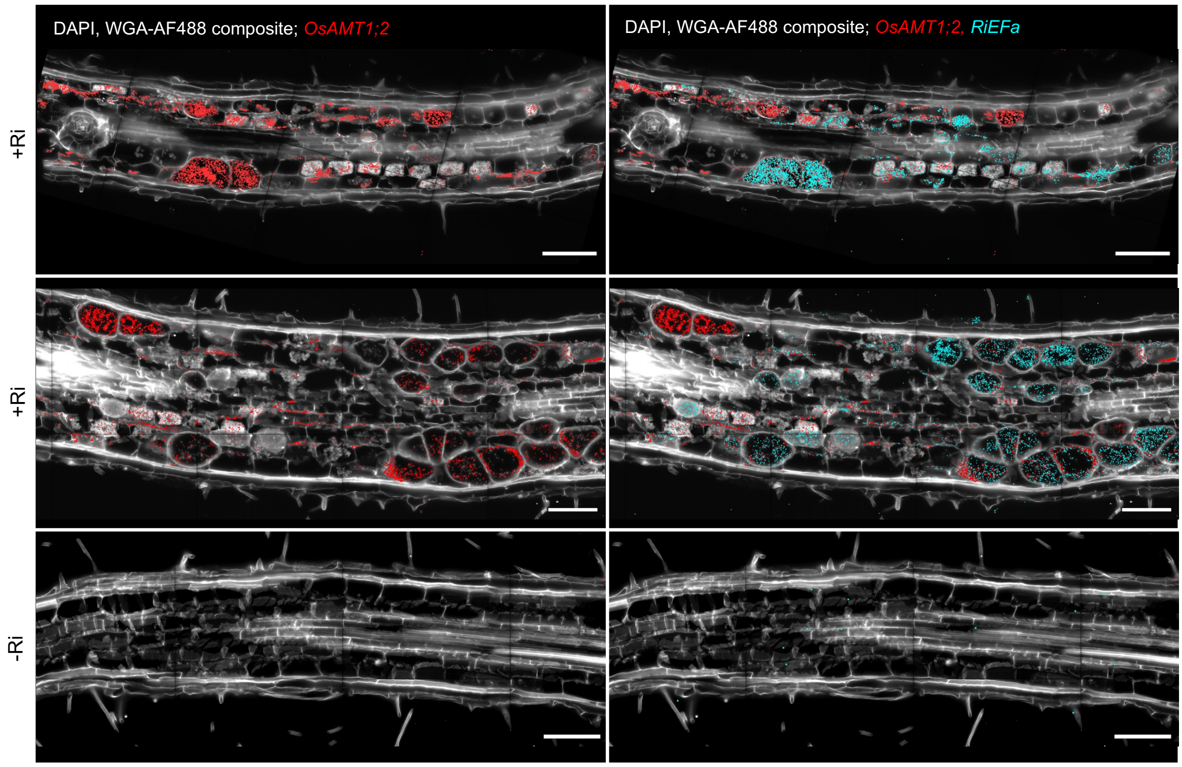


**Figure S10. Distribution of *OsAMT1;2* transcripts in mycorrhizal roots.** Image-transcript overlays highlighting spatial patterning of *OsAMT1;2* in +Ri and -Ri sections. DAPI/WGA-AF488 composite in white, each red spot corresponds to one detected transcript. Scale bar, 100 µm.

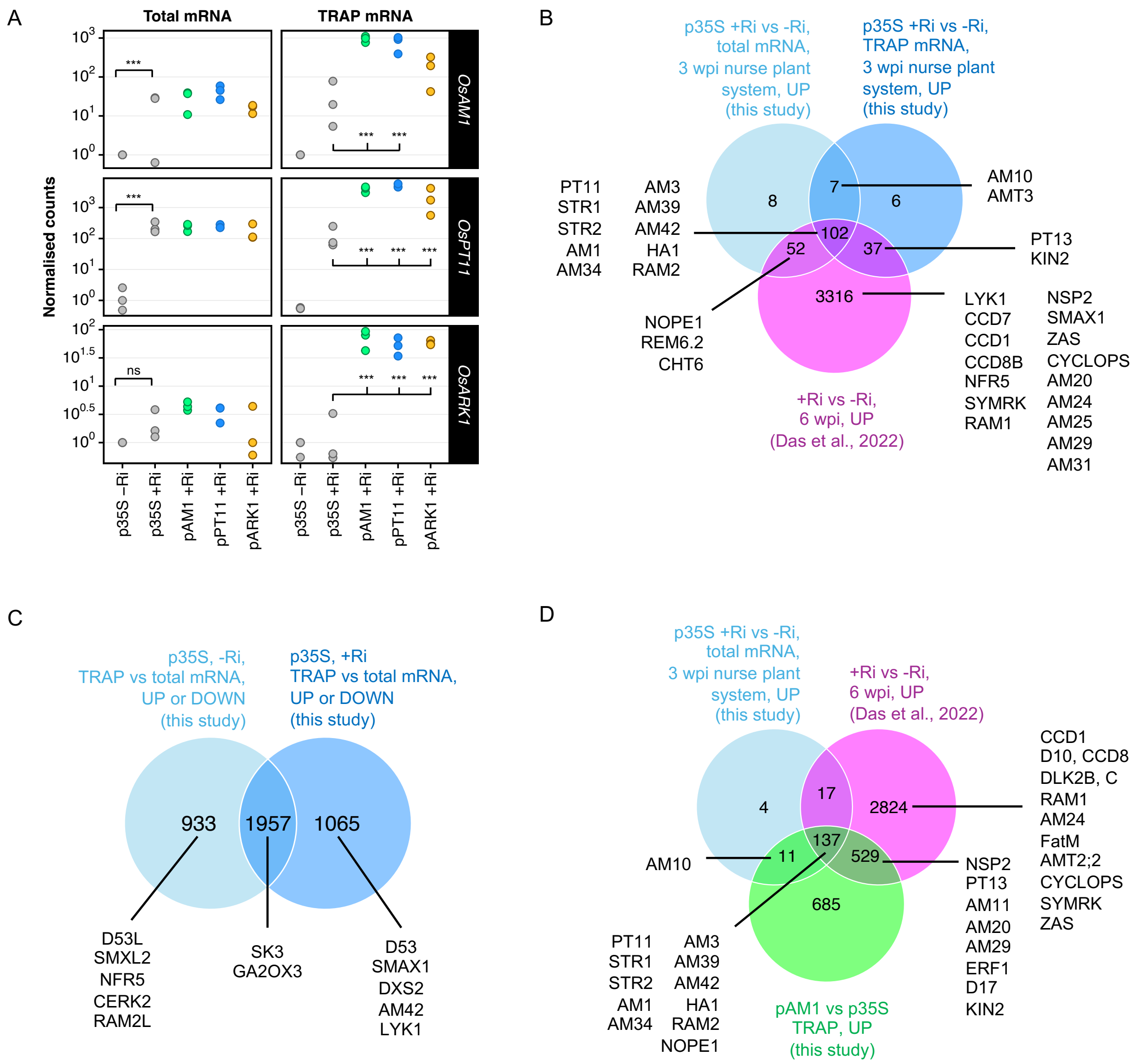


**Figure S11. Validation of AM-inducible TRAP-seq of -Ri and +Ri roots.** (A) Normalised gene counts for *OsAM1*, *OsPT11* and *OsARK1* in total and TRAP mRNA fractions for *p35S*, *pAM1*, *pPT11* and *pARK1:TRAP* lines. Significance levels as determined by DESeq2 shown with asterisks (* for p-value <0.05, ** for < 0.01, *** for < 0.001). (B) Venn diagram of overlaps between up-regulated genes for +Ri vs -Ri comparisons in this study for total and TRAP mRNA at three weeks post-inoculation (wpi) in a nurse plant set-up, with those from Das et al., 2022, at a later time-point (six wpi). Selected genes of interest highlighted. (C) Venn diagram of overlaps between enriched or depleted for TRAP vs total mRNA comparisons in non-inoculated (-Ri) or inoculated (+Ri) conditions. Selected genes of interest highlighted. (D) Venn diagram of overlaps between up-regulated genes for +Ri vs -Ri total mRNA comparisons in this study, those from Das et al., 2022, and enriched genes in *pAM1* vs *p35* TRAP mRNA. Selected genes of interest highlighted.

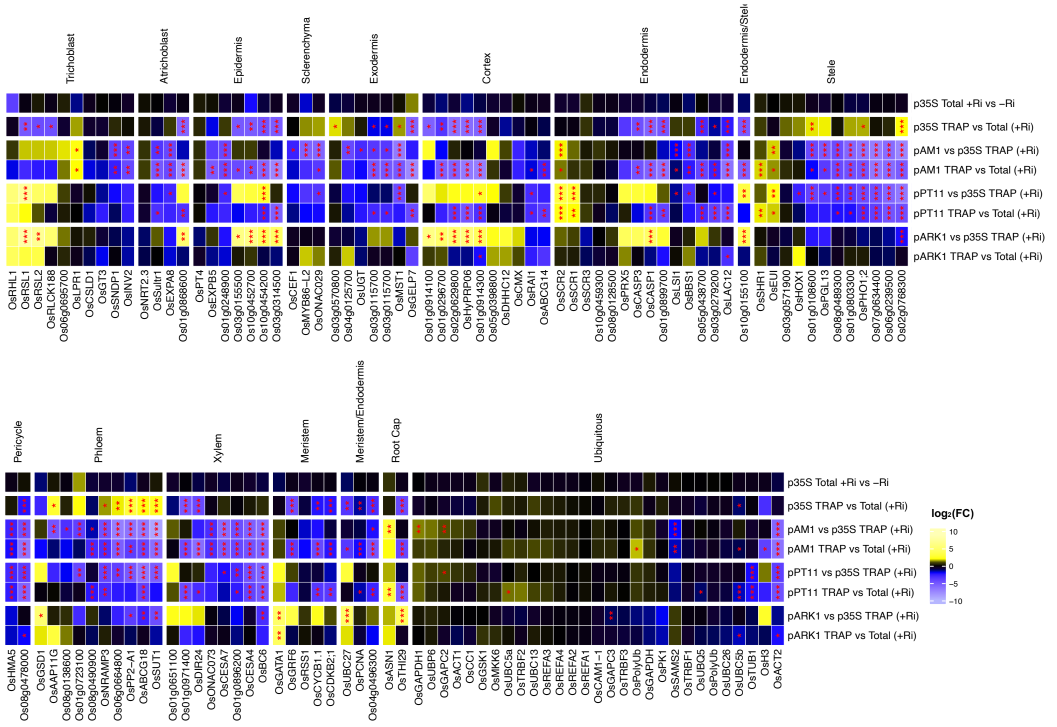


**Figure S12. Heatmap of $log_2FC$ for cell-identity markers**, predicted cell-type where expressed indicated above. Significance levels as determined by DGE using DESeq2 shown with asterisks (* for p-value <0.05, ** for < 0.01, *** for < 0.001). Genes were subjected to hierarchical clustering within each group.

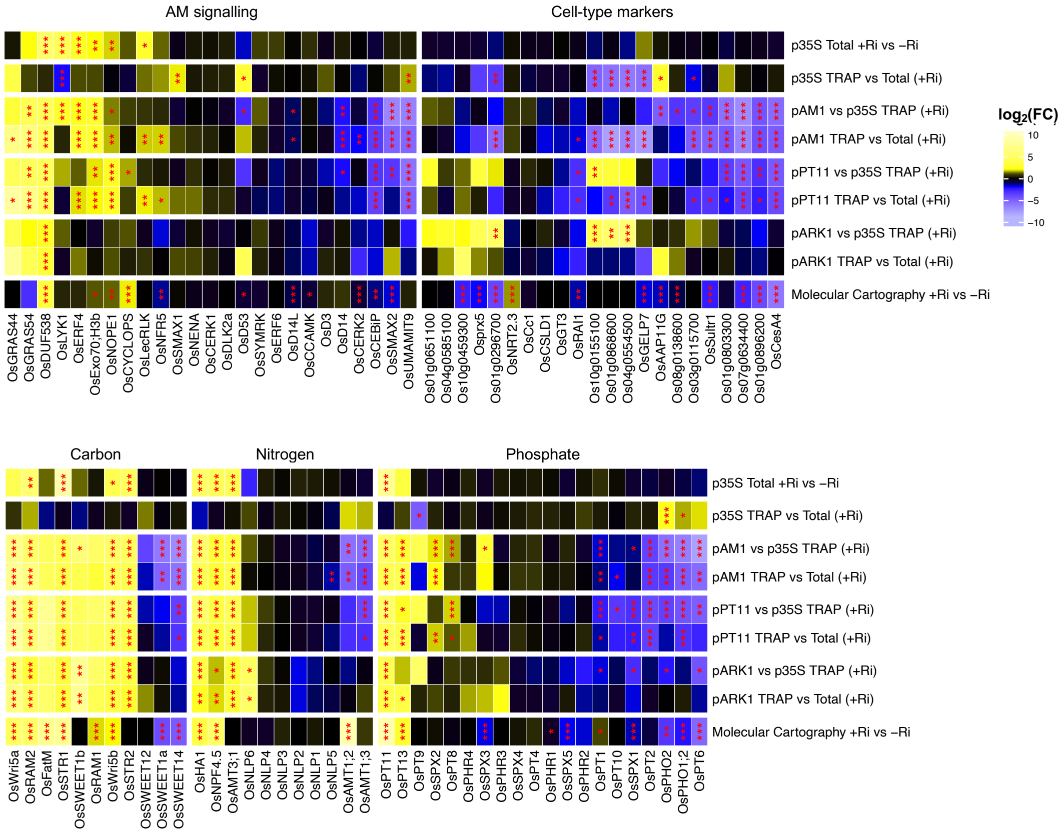


**Figure S13. Heatmap of $\log_2$FC for genes in the Molecular Cartography probe panel**, including fold-changes of the +Ri vs -Ri comparison of spatial transcriptomics transcript spot count data. Significance levels as determined by DGE using DESeq2 shown with asterisks (* for p-value <0.05, ** for < 0.01, *** for < 0.001). Genes were subjected to hierarchical clustering within each group.

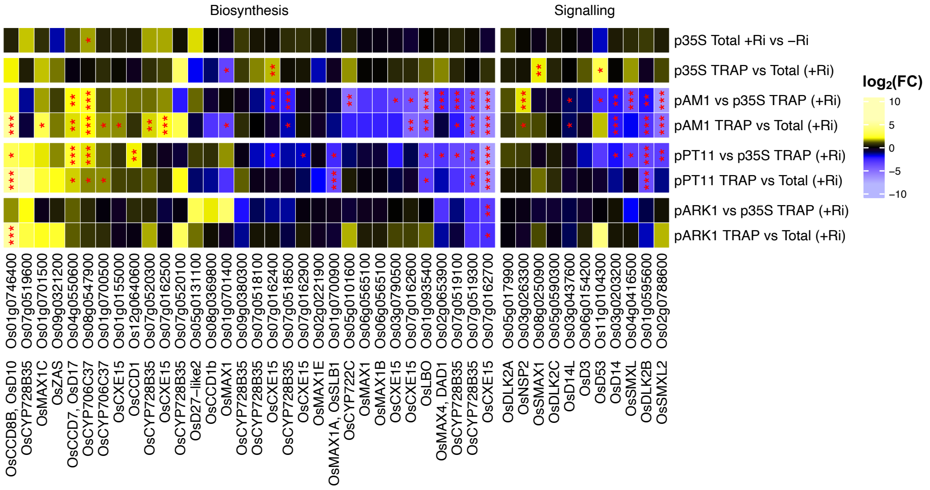


**Figure S14. Heatmap of $log_2$FC for genes involved in strigolactone/apocarotenoid biosynthesis**, perception and signalling. Significance levels as determined by DGE using DESeq2 shown with asterisks (* for p-value <0.05, ** for < 0.01, *** for < 0.001). Genes were subjected to hierarchical clustering within each group.

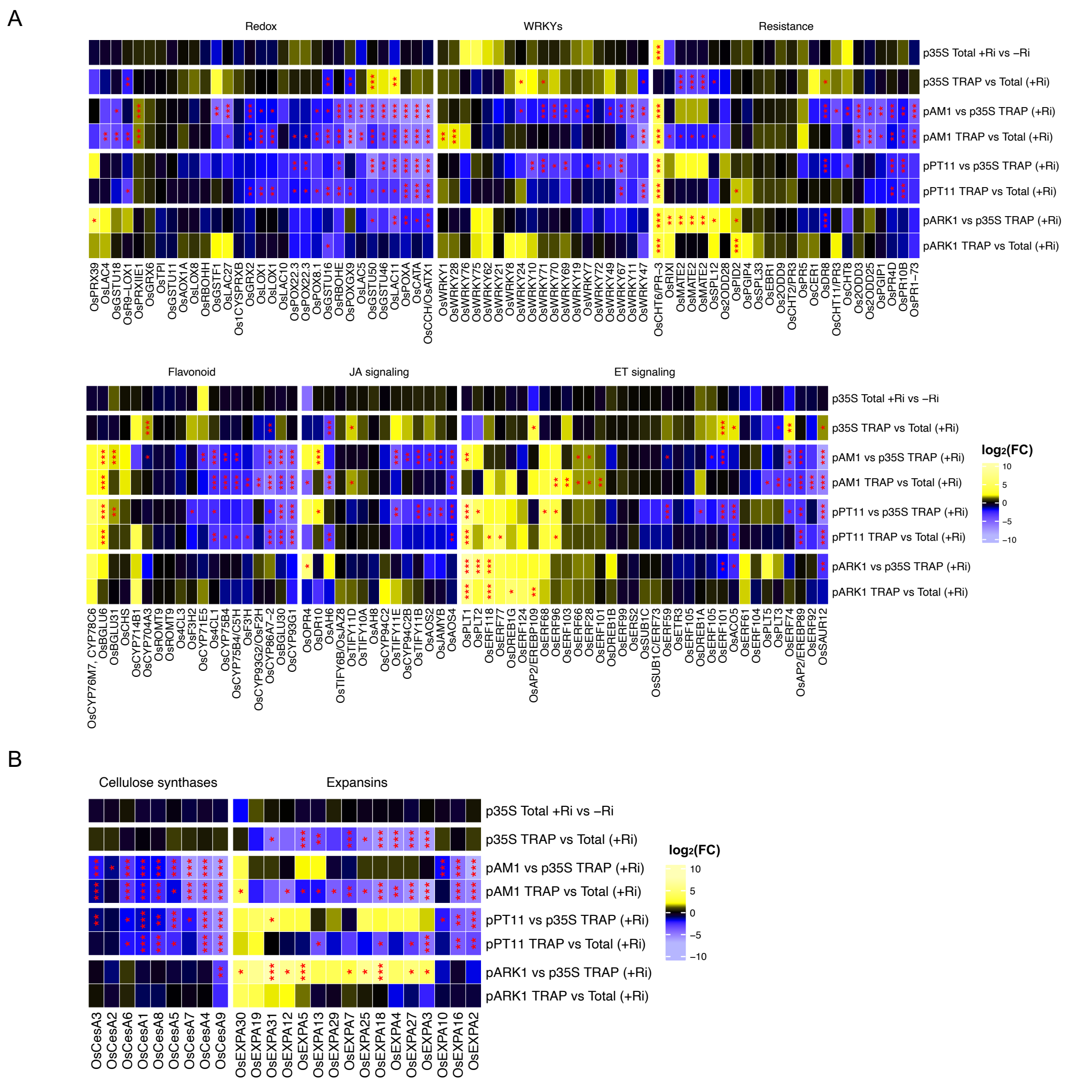

**Figure S15. Heatmap of $log_2FC$ for a selection of genes related to various processes (indicated above) associated with immunity and defence**, compiled by Tang et al., 2021(A), **cellulose synthases and expansins** (B). Significance levels as determined by DGE using DESeq2 shown with asterisks (* for p-value <0.05, ** for < 0.01, *** for < 0.001). Genes were subjected to hierarchical clustering within each group.

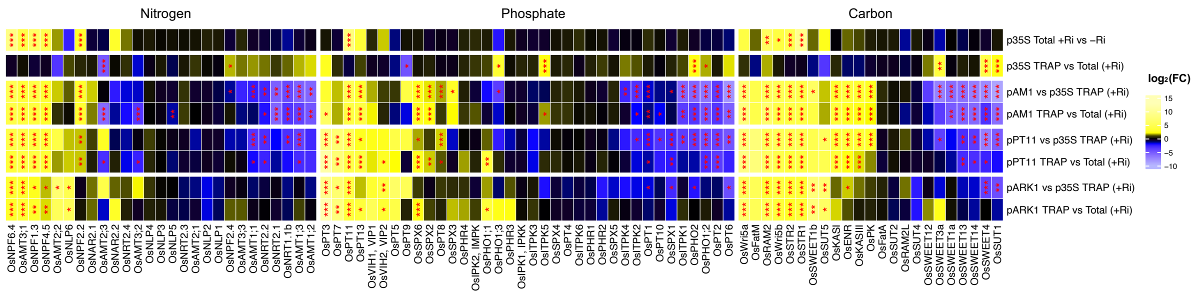


**Figure S16. Heatmap of $log_2$FC for key genes related to phosphate, nitrogen and carbon transport, perception and signalling**. Significance levels as determined by DGE using DESeq2 shown with asterisks (* for p-value <0.05, ** for < 0.01, *** for < 0.001). Genes were subjected to hierarchical clustering within each group.
